# Estrogen-related receptor beta activation and isoform shifting by cdc2-like kinase inhibition restricts migration and intracranial tumor growth in glioblastoma

**DOI:** 10.1101/558775

**Authors:** DM Tiek, SA Khatib, CJ Trepicchio, MM Heckler, SD Divekar, JN Sarkaria, E Glasgow, RB Riggins

## Abstract

Glioblastoma (GBM; grade 4 glioma) is a highly aggressive and incurable tumor. GBM has recently been characterized as highly dependent on alternative splicing, a critical driver of tumor heterogeneity and plasticity. Estrogen-related receptor beta (ERRβ, *ESRRB, NR3B2*) is an orphan nuclear receptor expressed in the brain, where alternative splicing of the 3’ end of the pre-mRNA leads to the production of three validated ERRβ protein products – ERRβ short form (ERRβsf), ERRβ2, and ERRβ exon 10-deleted (ERRβ-Δ10). Our prior studies have shown the ERRβ2 isoform to play a role in G2/M cell cycle arrest and induction of apoptosis, in contrast to the function of the shorter ERRβsf isoform in senescence and G1 cell cycle arrest. In this study, we sought to better define the role of the pro-apoptotic ERRβ2 isoform in GBM. We show that the ERRβ2 isoform is located in the nucleus, but also the cytoplasm. ERRβ2 suppresses GBM cell migration, interacts with the actin nucleation-promoting factor cortactin, and an ERRβ agonist is able to remodel the actin cytoskeleton and similarly suppress GBM cell migration. We further show that inhibition of the splicing regulatory cdc2-like kinases (CLKs) in combination with an ERRβ agonist shifts isoform expression in favor of ERRβ2 and potentiates inhibition of growth and migration in GBM cells and intracranial tumors.

## INTRODUCTION

Glioblastoma (GBM; grade 4 glioma) is a highly aggressive tumor, incurable, and markedly resistant to most systemic chemotherapies. Surgery, radiation, and adjuvant treatment with the DNA alkylating/methylating agent temozolomide (TMZ) are the current first-line standard of care. However, development of TMZ-resistance is rapid, which is in part due to re-expression of O6-methylguanine methyltransferase (MGMT), the DNA repair enzyme which removes TMZ-induced O6-methylguanine DNA adducts. Median survival is only ~14-16 months with TMZ, and after resistance has developed there is no established second-line regimen (1, 2). This is due in part to our limited understanding of the molecular drivers of GBM, and the inherent challenge of developing effective therapeutic strategies that penetrate the blood brain barrier (BBB). GBM is also a heterogeneous and invasive tumor that disseminates into healthy brain tissue. Even when the bulk of the tumor is surgically resected, chemotherapeutic options to target residual disease are limited by the BBB (3), which has severely limited repurposing of drugs from other cancers for GBM.

Dysregulation of alternative mRNA splicing is one potential mechanistic driver of GBM (4), given that the brain contains the most alternatively spliced transcripts of any organ and the expression of many splicing factors is upregulated in GBM vs. normal brain (5, 6). Serine/arginine rich (SR) proteins are a prominent group of splicing regulatory factors that are phosphorylated and thereby regulated by the cdc2-like kinases (CLKs) (7), some of which have been mechanistically implicated in GBM (8). While CLK inhibitors have not yet entered clinical trials, preclinical studies of TG-003 (a pan-CLK inhibitor) show that this agent can cross the BBB in mouse models of autism (9). Ongoing clinical trials are testing first-generation splicing regulatory drugs, such as H3B 8800 for myelodysplastic syndromes, acute myeloid leukemia, and chronic myelomonocytic leukemia (10).

Given that improved therapeutic options are an urgent clinical need for GBM, the nuclear receptor superfamily - members of which are highly successful targets in breast and prostate cancers - provide another novel target strategy. Estrogen-related receptor beta (ERRβ, *ESRRB, NR3B2*) is the founding orphan member of the nuclear receptor superfamily (11). By definition, orphan nuclear receptors lack known endogenous ligands, though their function can be modified by coregulatory proteins or synthetic ligands that increase or decrease their transcription factor activity (12, 13). DY131 is one synthetic agonist that has been shown to enhance ERRβ transcription factor activity in the murine arcuate nucleus (14, 15), which is strongly suggestive of BBB penetrance. ERRβ is alternatively spliced at the 3’ end, leading to the production of ERRβ short form (ERRβsf), ERRβ2, and ERRβ exon 10-deleted (ERRβ-Δ10) (Figure 1A, (16)). These 3’ splicing events are unique to primates, with all lower vertebrate organisms containing genomic sequences for only the ERRβsf isoform (16). Inclusion of additional 3’ exons in ERRβ2 and ERRβ-Δ10 produces 67 and 75 amino acid carboxyl terminal extensions, or F domains, which can modify transcription factor function and recruit distinct coregulatory proteins (17).

**Figure 1.**
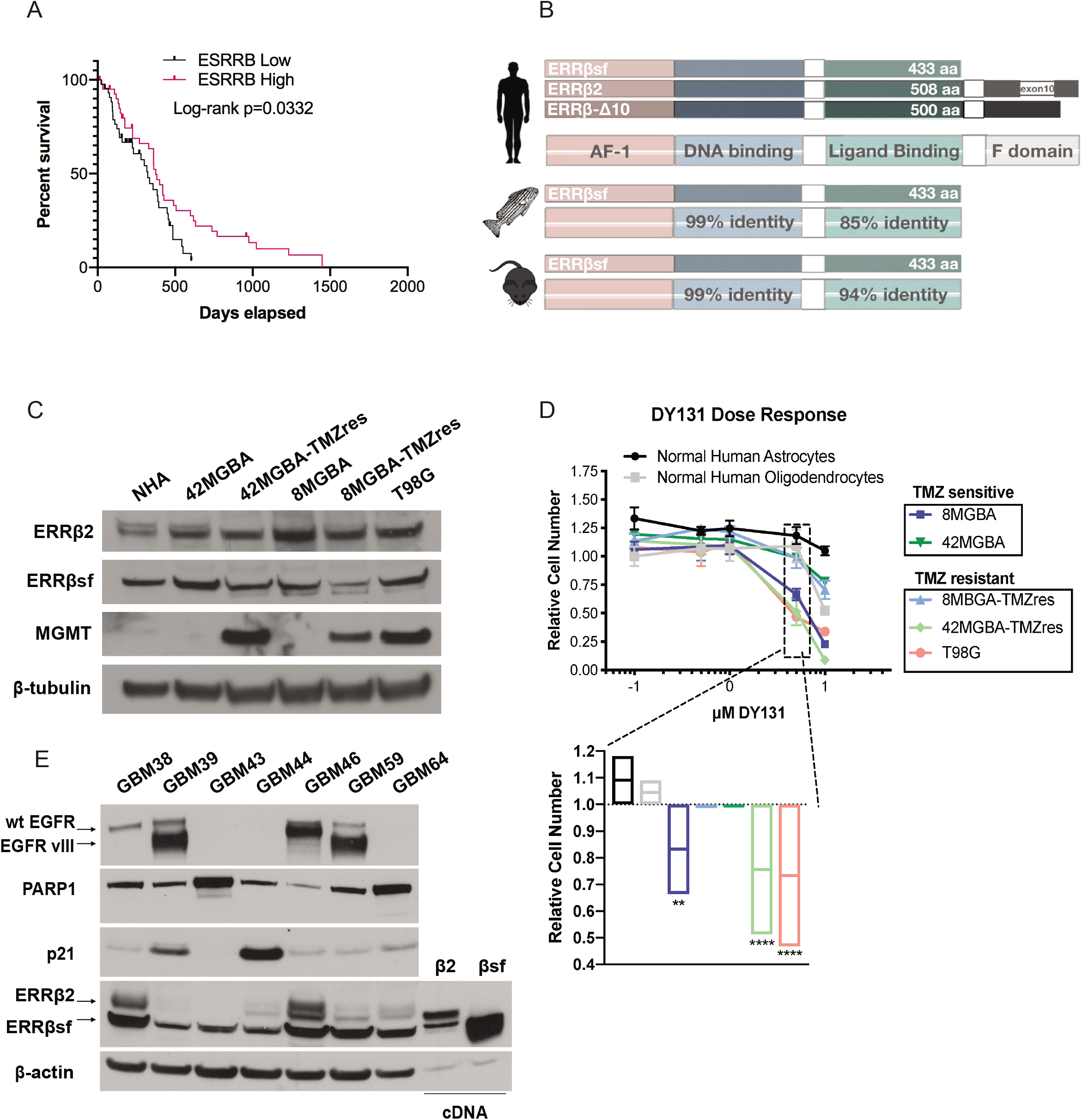
**(A)** Expression of ESRRB correlates with GBM survival. Normalized RNAseq data (Fragments Per Kilobase of transcript per Million mapped reads upper quartile, FPKM-UQ) from the GBM TCGA dataset were downloaded from XenaBrowser. Log-rank survival analysis was performed for upper vs. lower quartile of ESRRB expression, p=0.0332. **(B)** Differential splicing at the 3’ end of the *ESRRB* pre-mRNA leads to the production of three known ERRβ transcripts and protein products—the short form (ERRβsf), and two longer forms called beta2 (ERRβ2), and exon 10-deleted (Δ10). The ERRβsf isoform is conserved in zebrafish and mice, with percent identity for each ortholog compared to the human sequence. AF-1 = activation function-1, aa = amino acid. **(C)** ERRβ2 and ERRβsf protein are expressed in primary normal human astrocytes (NHAs) as well as in multiple TMZ sensitive (8MGBA, 42MGBA) and resistant (8MGBA-TMZres, 42MGBA-TMZres, T98G) GBM cell lines. O6-methylguanine methyltransferase (MGMT), a known marker of therapy resistance in GBM, is heavily expressed in the resistant cell lines. **(D)** DY131, a small molecule agonist of ERRβ, inhibits cellular proliferation in TMZ sensitive and resistant cell lines but not in NHAs or normal human oligodendrocytes. Data in the growth curves are presented as the mean (point) ± standard deviation (SD, error bars), and data in the inset are presented as the mean (line) ± minimum/maximum values (box) for 3 (NHA), 6 (oligodendrocytes, T98G), or 8 (all other cell lines) technical replicates, which were analyzed by two-way analysis of variance (ANOVA) with *post hoc* Dunnett’s multiple comparisons test. Data are representative of at least two independent biological replicates. **, *** denote p<0.01 and p<0.0001, respectively, vs. DMSO control. **(E)** Expression of ERRβ2 and ERRβsf vary across multiple patient-derived xenografts (PDXs) of GBM, as does wild type (wt) and vIII mutant EGFR, the DNA damage marker PARP, and the cyclin-dependent kinase inhibitor, p21. Lanes labeled β2 and βsf denote lysate from cells transfected with the indicated cDNA construct.

We previously identified the ERRβ2 isoform as a cytoplasmic and centrosome-adjacent protein (18), and showed that activation of this isoform can delay mitosis and partially repress the transcription factor activity of the ERRβsf isoform (19). The goal of this study was to better define the function and interacting partners of the pro-apoptotic ERRβ2 isoform in GBM and test the ability of splicing kinase inhibitors to shift isoform balance towards the ERRβ2 isoform. We found that the cytoplasmic ERRβ2 isoform suppresses GBM cell migration and interacts with the actin nucleation-promoting factor cortactin, and that an ERRβ agonist remodels the actin cytoskeleton and also suppresses migration. CLK inhibition with TG-003 in combination with the ERRβ agonist DY131 shifts isoform expression in favor of ERRβ2 and leads to suppression of growth and migration in TMZ-resistant GBM cells. Finally, we use a novel zebrafish model to show that the combination of TG-003 and DY131 has anti-tumor activity *in vivo* in a setting where the BBB is intact.

## METHODS

### Cell Lines and Culturing Conditions

Primary normal human astrocytes (NHAs) were purchased from Lonza (#CC-2565). Immortalized human oligodendrocyte MO3.13 cells were a kind gift from Dr. Alexandra Taraboletti (Lombardi Comprehensive Cancer Center, LCCC). Temozolomide (TMZ) sensitive 42MGBA and 8MGBA cell lines were provided by Dr. Jeffrey Toretsky (LCCC), and the *de novo* TMZ resistant T98G cell line was provided by Dr. Todd Waldman (LCCC). Acquired TMZ resistant 42MGBA-TMZres and 8MGBA-TMZres cell line variants were developed by our lab and previously described (20). All cells tested negative for *Mycoplasma* contamination and were maintained in a humidified incubator with 95% air: 5% carbon dioxide. All cell lines were fingerprinted by the LCCC Tissue Culture Shared Resource to verify their authenticity using the standard 9 STR loci and Y-specific amelogenin. Both the 42MGBA-TMZres and 8MGBA-TMZres are documented to be of the same origin as their respective parental cell lines. NHAs were used within one passage and maintained in astrocyte growth medium (AGM, Lonza #CC-3187) supplemented with L-glutamine, gentamicin sulfate, ascorbic acid, human epidermal growth factor (HEGF), insulin and 3% fetal bovine serum (FBS) (Lonza, #CC-4123). MO3.13, 42MGBA, 8MGBA, 42MGBA-TMZres, and T98G cells were grown in Dulbecco’s Modified Eagle Medium (DMEM, high glucose, ThermoFisher, #11965092) with 10% FBS. 8MGBA-TMZres cells were grown in DMEM with 10% FBS and 100 μM TMZ. TMZ (Selleckchem, #S1237) was dissolved in dimethyl sulfide (DMSO, Sigma, #D8418) to 130 mM and used at concentrations indicated. DY131 (Tocris, #2266) was dissolved in DMSO to 10mM and used at the concentrations indicated.

### Western Blotting

Cells were lysed in RIPA buffer supplemented with protease and phosphatase inhibitors (Roche, #4906837001) for protein extractions and separated by polyacrylamide gel electrophoresis using 4-12% gradient gels (Novex by Life Tech, #NP0321BOX) as described previously (19). They were then transferred onto Nitrocellulose membranes (Invitrogen, #IB23001) with the iBlot2 (Invitrogen, #IB21001) and probed with the following antibodies: ERRβ2 (1:500, RNDSystems, #PP-H6707-00), ERRβsf (1:1000, RNDSystems, #PP-H6705-00), PARP (1:1000, Cell Signaling, #9542L), CDKN1A/p21 (1:300, Santa Cruz Biotechnology, #sc-756), EGFR (1:1000, Cell Signaling, #2232S), MGMT (1:1000, Cell Signaling, #2739S), Cortactin (1:1000, Upstate, #05-180), phosphorylated SR proteins (clone 1H4, 1:500, Millipore, #MABE50), total SR proteins (1;1000, Sigma-Aldrich, #MABE126), phosphorylated histone H3 Serine 10 (1:1000, Cell Signaling, #3377S), and total histone H3 (1:1000, Cell Signaling, #9715S). Beta-Tubulin (1:5000, Sigma Aldrich, #T7816) and Beta-Actin (1:5000, Sigma-Aldrich, #A5316) were used as a loading controls. Proteins were detected with horseradish peroxidase-conjugated secondary antibodies (1:5000, GE Healthcare Life Sciences, #NA931-1ML (Mouse) or #NA934-1ML (Rabbit)) and enhanced chemiluminescent detection HyGLO Quick Spray Chemiluminescent (Denville Scientific, #E2400) using film (Denville Scientific, #E3212).

### Immunofluorescence

Cells were seeded at a density of 40,000-50,000 cells onto 18mm diameter #1.5 round coverslips (VWR, #101413-518) in 12-well dishes. On the following day, the media was removed and cells were fixed and permeabilized in 3.2% paraformaldehyde (PFA) with 0.2% Triton X-100 in PBS for 5 minutes at room temperature. Three washes were performed with PBS in the 12-well plate, then coverslips were inverted onto 120 μL of primary antibody in the antibody block (0.1% gelatin with 10% normal donkey serum in PBS) on strips of parafilm and incubated for one hour. Coverslips were first incubated with either ERRβ2 (1:150) or ERRβsf (1:200) for 1 hour. After incubation with primary antibodies, coverslips were washed three times with PBS. Then coverslips were inverted onto 100 μL of antibody block with secondary antibodies (Alexa Fluor 488 anti-mouse - 1:200, Life Technologies #A11029) and DAPI (DNA, 1:500 dilution) for 20 minutes in the dark. Coverslips were again washed 3x with PBS, then gently dipped four times into molecular biology-grade water before inversion onto one drop of Fluoro-Gel (with TES Buffer, Electron Microscopy Sciences, #17985-30) then allowed to air-dry in the dark for at least 10 minutes. Slides were stored at 4°C until image collection on the LCCC Microscopy & Imaging Shared Resource’s Leica SP8 microscope with the 63X oil objective.

### Cell Growth Assays

Cells were seeded in 96-well plastic tissue culture plates at 1000 cells/well one day prior to treatment with the indicated concentrations of DY131. Cells were treated for a total of 8-10 days, with media changed and drug replenished on Days 4 or 5. Staining with crystal violet, resolubilization, and analysis of staining intensity as a proxy for cell number was performed as described previously (19).

### <u>I</u>mmunohistochemistry on Human GBM Tumor Samples

Immunohistochemical staining of GBM human tumor samples was performed for ERRβ2 or ERRβsf. Five micron sections from formalin-fixed paraffin-embedded tissues were de-paraffinized with xylenes and rehydrated through a graded alcohol series.

For ERRβ2, heat induced epitope retrieval (HIER) was performed by immersing the tissue sections at 98°C for 20 minutes in 10 mM citrate buffer (pH 6.0) with 0.05% Tween. Immunohistochemical staining was performed using a horseradish peroxidase labeled polymer from Agilent (#K4001, #K4003) according to manufacturer’s instructions. Briefly, slides were treated with 3% hydrogen peroxide and 10% normal goat serum for 10 minutes each, and exposed to primary antibody at a 1:125 dilution overnight at 4°C. Slides were exposed to the appropriate HRP labeled polymer for 30min and DAB chromagen (Dako) for 5 minutes.

For ERRβsf, HIER was performed by immersing the tissue sections at 98oC for 20 minutes in 110 mM Tris, 1m M EDTA pH 9.0 buffer (Genemed). Immunohistochemical staining was performed using the VectaStain Kit from Vector Labs according to manufacturer’s instructions. Briefly, slides were treated with 3% hydrogen peroxide, avidin/biotin blocking, and 10% normal goat serum and exposed to primary antibody at a 1:200 dilution overnight at 4°C. Slides were exposed to appropriate biotin-conjugated secondary antibodies (Vector Labs), Vectastain ABC reagent and DAB chromagen (Dako).

For both primary antibodies, slides were counterstained with Hematoxylin (Fisher, Harris Modified Hematoxylin), blued in 1% ammonium hydroxide, dehydrated, and mounted with Acrymount. Consecutive sections with the primary antibody omitted were used as negative controls.

### ERRβ Knockdown

Short hairpin RNA (shRNA)-mediated silencing of ERRβ2 and ERRβsf in T98G GBM cells was reported previously (19). shRNA sequences targeting ESRRB are as follows: shESRRB-1: TGAGGACTACATCATGGAT shESRRB-2: TGCAGCACTTCTATAGCGT.

### Migration Assays

For scratch wound assays, cells were plated at 150,000-200,000 cells/well depending on the cell line and allowed 48 hours to create a monolayer. After monolayer formation, a P200 tip was used to make a scratch and images were taken at 0 hr, 24 hr, 48 hr, and 72 hr time points. Analysis was done in ImageJ (21) to determine percent closed with 0% being at 0hr. For transwell migration assays, cells were cultured in low serum conditions (0.5% FBS) overnight. The following day, 100,000 cells in 200 μL of 0.5% DMEM were loaded into the top of a modified Boyden chamber (24-well, 8 μm cell culture insert, Becton Dickinson #353097), which was then placed in a 24-well plate containing 500 μL of DMEM supplemented with full serum (10% FBS). Cells were allowed to migrate for 4 and 24 hours at 37°C, at which time the nonmigratory cells were removed from the top of the membrane with cotton swabs and the bottom of the membrane was stained with crystal violet as described for cell growth assays. After staining the membranes were excised from the insert, mounted on glass slides using Fluoro-Gel, and allowed to air dry. Cells per field at 20X magnification were counted by light microscopy.

### Immunoprecipitation

On day 0, cells were seeded at 200,000/well in 6-well dishes. The next morning on day 1, two separate tubes were made and then combined for the transfection. Tube #1 contained 2.5 μg plasmid DNA (psg5, ERRβ2, ERRβsf, or ERRβ-Δ10), 2.5 μL PLUS reagent (from Lipofectamine LTX, Life Technologies, #15338100), and Opti-MEM (Life Technologies, #31985070) that brings the total volume to 50 μL. Tube #2 contained 500 μL Opti-MEM and 6.25 μL Lipofectamine LTX. Tube #1 was then added to tube #2 and pipetted up and down a few times to mix thoroughly. The now tube #2 with all reagents was left untouched for 25-30 minutes. Before transfecting, medium was changed to serum free DMEM. Then the transfection reagent was slowly taken up in a P1000 tip and added drop-wise to the cells. 4-6 hours later, medium was changed back to 10% DMEM overnight. The next morning (day 2), medium was changed again with fresh 10% DMEM. In the afternoon of day 2, cells were lysed in RIPA for protein extraction. Protein was quantified, and amounts were normalized between all samples to be used in the immunoprecipitation (IP). Total volume for each IP was brought up to 500 μL of RIPA with inhibitors. An aliquot was also set aside for input for each sample. In the evening of day 2, 1 μL of the cortactin antibody (Abcam, #ab81208 - 1:500 dilution) was added to each sample and allowed to rotate overnight at 4°C. The next morning, Protein A/G beads (Pierce, #20421) were vortexed, and 30 μL of beads were added to an eppendorf tube per sample. Beads were then washed in cold RIPA buffer with inhibitors and centrifuged at 4°C for 5 minutes. Supernatant was aspirated, and cold RIPA buffer with inhibitors was added to bring up the volume. Then 30 μL of washed beads were added to each IP tube and allowed to rotate again at 4°C for 1 hour. Tubes were then centrifuged for 5 minutes at 10,000 rpm at 4°C. Supernatant was aspirated, and 500 μL of cold RIPA with inhibitors was used to wash IPs. Tubes were then centrifuged again for 5 minutes at 10,000 rpm at 4°C. IPs were washed 2x more in tris-saline buffer (50 mM tris base pH 7.5, 150 mM NaCl in diH_2_O). After the last wash, aspirate supernatant and add 30 μL of 2X loading buffer. Samples were then vortexed and boiled for 9 minutes. Samples were then spun down in a centrifuge at room temperature for 1-2 minutes. Finally, both IP and input samples were loaded onto a 4-12% gradient gel and followed the Western Blotting protocol from above.

### F-actin Flow Cytometry

Cells were seeded at 100,000 - 200,000 cells/well in 6-well plastic tissue culture dishes then treated with 5 μM DY131 the next day. After 24 hours, cells were collected by trypsinization, combined with nonadherent cells in the culture media, then washed with PBS prior to fixation in 4% PFA for 5 minutes. Fixed cells were permeabilized using 0.2% Triton X-100 in PBS for 5 minutes, then blocked in 0.1% gelatin with 10% normal donkey serum in PBS for 30 minutes at room temperature. Cells were stained in suspension with a 1:300 dilution of Anti-Stain 488 Fluorescent Phalloidin (Cytoskeleton, #PHDG1), gently mixed, and incubated in the dark at room temperature for 30 minutes. Stained cells were washed twice with PBS, then 30,000 cells were acquired by flow cytometry on a BectonDickinson Fortessa. Data was analyzed using FCSExpress 6 (DeNovo Software, Glendale, CA).

### Cell Cycle Analysis

On day 0, cells were seeded at 200,000 cells per well in 6-well plastic tissue culture dishes one day prior to treatment with the indicated concentrations of drug. For experiments with TG-003 +/− DY-131, treatment time was 24 hours. After 24 hours, cells were collected, washed with PBS, ethanol-fixed, stained with propidium iodide, and analyzed for cell subG1 (fragmented/apoptotic) DNA content and cell cycle profile. 30,000 cells were acquired by flow cytometry on a BectonDickinson Fortessa. Files were modeled using ModFit software (Verity Software, Topsham, ME) to determine subG1, G1, S, and G2/M cell cycle stage.

### Zebrafish Intracranial Xenografts

The zebrafish model used is a transgenic line with green blood vessels in a double mutant, transparent, background with the genotype Tg(kdrl:GRCFP)zn1; mitfab692/ b692; ednrb1b140/ b140. This line is propagated in house by raising equal numbers of fry from five independent group matings, each consisting of three females and three males, in order to maintain background genetic diversity. Correct genotype is determined by visual inspection; the fish lack all pigment except for the eyes and the blood vessels fluoresce bright green. A new generation is raised bi-yearly and the oldest generation is discontinued when they reach two years of age. All procedures were performed in accordance with NIH guidelines on the care and use of animals and were approved by the Georgetown University Institutional Animal Care and Use Committee, Protocol #2017-0078.

Tumor cells were labeled in suspension with a 1:100 dilution of Vibrant CM-DiI cell-labeling solution (ThermoFisher, #V22885) at a density of 5 x 10^6^ cells/ml for 20 min at 37°C. The labeled cells were washed 5 times in PBS, suspended in PBS at 5 x 10^7^ cells/ml, and then backloaded into a pulled borosilicate microinjection needle (David Kolf Instruments). The needles were placed in a vertical position with the point facing down for 10 min to allow the cells to settle toward the tip. Thirty-six (36) hours post fertilization (hpf) embryos were anesthetized in 160 μg/ml buffered tricaine solution (Sigma-Aldrich). The embryos were loaded onto an injection plate at 37°C and covered with 1.5% low melting point agarose (Fisher Scientific) where they were positioned for injection. The plate was allowed to cool to room temperature and then was covered in fish water (0.3 g/L sea salt) containing 160 mg/ml tricaine. The embryos were injected with 1-2 nl of cell suspension (50-100 cells) into the area of the midbrain-hindbrain border. Following injection, the embryos were manually freed from the agarose and allowed to recover at 28°C for one hour, followed by incubation at 33°C. At 5 days post fertilization (dpf), tricaine-anesthetized embryos were mounted in 3% methyl cellulose and imaged on an Olympus XI-71 Inverted epifluoresce microscope. Following imaging, the fry were transferred to 24-well plates in 1ml fish water per well. At 10 dpf, fry were anesthetized with 112 μg/ml tricaine, mounted in 3% methyl cellulose, and imaged on an Olympus XI-71 Inverted epifluoresce microscope, or, mounted in 1.5% low melting point agarose for imaging on a Leica SP8 confocal microscope.

Treatment groups were assigned to achieve similar numbers of larvae containing similar distributions of tumor sizes. The larvae were treated starting at 6 dpf. Final drug concentrations were made using 1:1000 dilution of stock in DMSO. Control larvae were treated with 0.1% DMSO. The drugs were refreshed daily by removing most of the solution, leaving ~100 μl to keep the fry wet, followed by the addition of 1 ml of fresh drug solution. For blood-brain barrier (BBB) imaging experiments, Cascade Blue dextran, 10,000 MW (ThermoFisher, #D1976) at 25 mg/ml was back-loaded into a pulled borosilicate microinjection needle. 10 dpf larvae were anesthetized in 112 μg/ml buffered tricaine solution and loaded onto an injection plate at 37°C and covered with 1.5% low melting point agarose, where they were positioned for injection. Approximately 2 nl of Cascade Blue dextran solution was injected into the common cardinal vein. The injected larvae were then mounted in 1.5% low melting point agarose and imaged on a Leica SP8 confocal microscope at 1 hour post injection.

### Computational, Image, and Statistical Analysis

Images and figures were compiled using either Adobe Photoshop or Illustrator. Densitometry of the ERRβ2:ERRβsf ratio was calculated using ImageJ (21). Calculation of zebrafish intracranial xenograft tumor size was carried out in Illustrator as follows: Zebrafish images were imported into Illustrator. The red channel was selected, which highlighted the GBM cells as DiI fluoresces red. The “magic wand” tool was chosen to select the red fluorescent tumor, where after selection area was calculated by Illustrator’s “measure area” function. These measurements were exported and pre- and post-treatment tumor areas were compared. Motif analysis of ERRβ2 F domain sequences was performed using ScanSite3.0 (22). Statistical analysis and graphing were performed using GraphPad Prism 8.0, with the following exceptions. Supplementary Figure 1A was generated using RNAseq data from EMBL-EBI (dataset E-MTAB-4840, (23, 24)). Figure 1A was generated using RNAseq data from XenaBrowser (25). Supplementary Figure 3A was generated using RNAseq data from the Chinese Glioma Genome Atlas through the International Cancer Genome Consortium (https://icgc.org) analyzed in R using GGally and ggplot2 packages. Supplementary Figure 3B was generated and analyzed in GraphPad Prism 8.0 from RNAseq data obtained through the Ivy Glioblastoma Atlas Project (26). SRSF6 consensus binding sites (27) in *ESRRB* exon 10 were predicted using RBPmap (28). Data are presented as the mean ± standard deviation (SD) unless otherwise indicated. The number of biological, technical replicates, and specific statistical tests performed are reported in the figure legend for each figure. Statistical significance is defined as α ≤ 0.05.

## RESULTS

The conserved ERRβsf and primate-specific ERRβ2 isoforms of ERRβ have distinct functions in cell cycle regulation, with activation of ERRβ2 by the synthetic agonist ligand DY131 causing mitotic arrest and apoptosis (19). These mechanistic discoveries demonstrated the ERRβ2 isoform to play a more anti-tumorigenic role than ERRβsf. Therefore, the goal of this study was to better define the function and interacting partners of the pro-apoptotic ERRβ2 isoform in GBM and test the ability of splicing kinase inhibitors to shift isoform balance towards the ERRβ2 isoform.

### ERRβ isoforms are expressed in GBM

Nuclear receptors are attractive drug targets, and successful inhibitors to nuclear receptors like estrogen receptor alpha (ERα) have fundamentally changed cancer treatment outcomes for breast cancer patients. However, targeting ERα with tamoxifen in GBM has been unsuccessful (29), in part because its expression in the brain is quite low, in contrast to estrogen-related receptor beta (gene symbol ESRRB), an orphan nuclear receptor expressed in the brain (**Supplementary Figure 1A**). In The Cancer Genome Atlas (TCGA), high (upper quartile) *ESRRB* mRNA expression is significantly associated with longer overall survival (**Figure 1A**). Two ERRβ protein isoforms - ERRβ2 and ERRβsf - are expressed across multiple TMZ sensitive and resistant GBM cell lines (20) and primary normal human astrocytes (NHAs, **Figure 1B,C**), as well as immortalized human oligodendrocytes (MO3.13, **Supplementary Figure 1B**). DY131 is a small molecule agonist of ERRβ that has anti-proliferative activity in several preclinical cancer models (18, 30). DY131 is growth inhibitory in GBM cell lines, but not NHAs or immortalized oligodendrocytes where it shows a slight increase in growth (**Figure 1D**). A limitation of conventional GBM cell lines is that they do not always recapitulate essential GBM molecular features or pathobiology (31). Patient-derived xenografts (PDXs) from primary tumors have been established to address this limitation, and we show that ERRβ2 and ERRβsf protein are broadly expressed across PDX specimens representative of multiple GBM molecular subtypes and clinical/pathological features (**Figure 1E, Supplementary Table 1**). EGFR wild type- and vIII-mutant amplified tumors are more frequently categorized in the Classical molecular subtype (32), while PARP1 expression is enriched in Classical and Proneural subtypes (33), and *CDKN1A*/p21 expression is indicative of primary vs. secondary GBM (34).

Studies in COS-1 cells transfected with ERRβ isoform cDNA show that ERRβsf localizes to the nucleus, while ERRβ2 is localized to the cytosol and nucleus, suggesting differential functions (16). This pattern of subcellular localization of endogenous ERRβsf (nucleus) vs. ERRβ2 (cytosol and nucleus) is similarly observed in NHAs, oligodendrocytes, and GBM cells (**Figure 2A, Supplementary Figure 1C**), and in primary tumor specimens (**Figure 2B**). Collectively, these data demonstrate that ERRβ isoforms are expressed, and a well-established ERRβ agonist is growth-inhibitory, in multiple TMZ sensitive and resistant GBM models.

**Figure 2.**
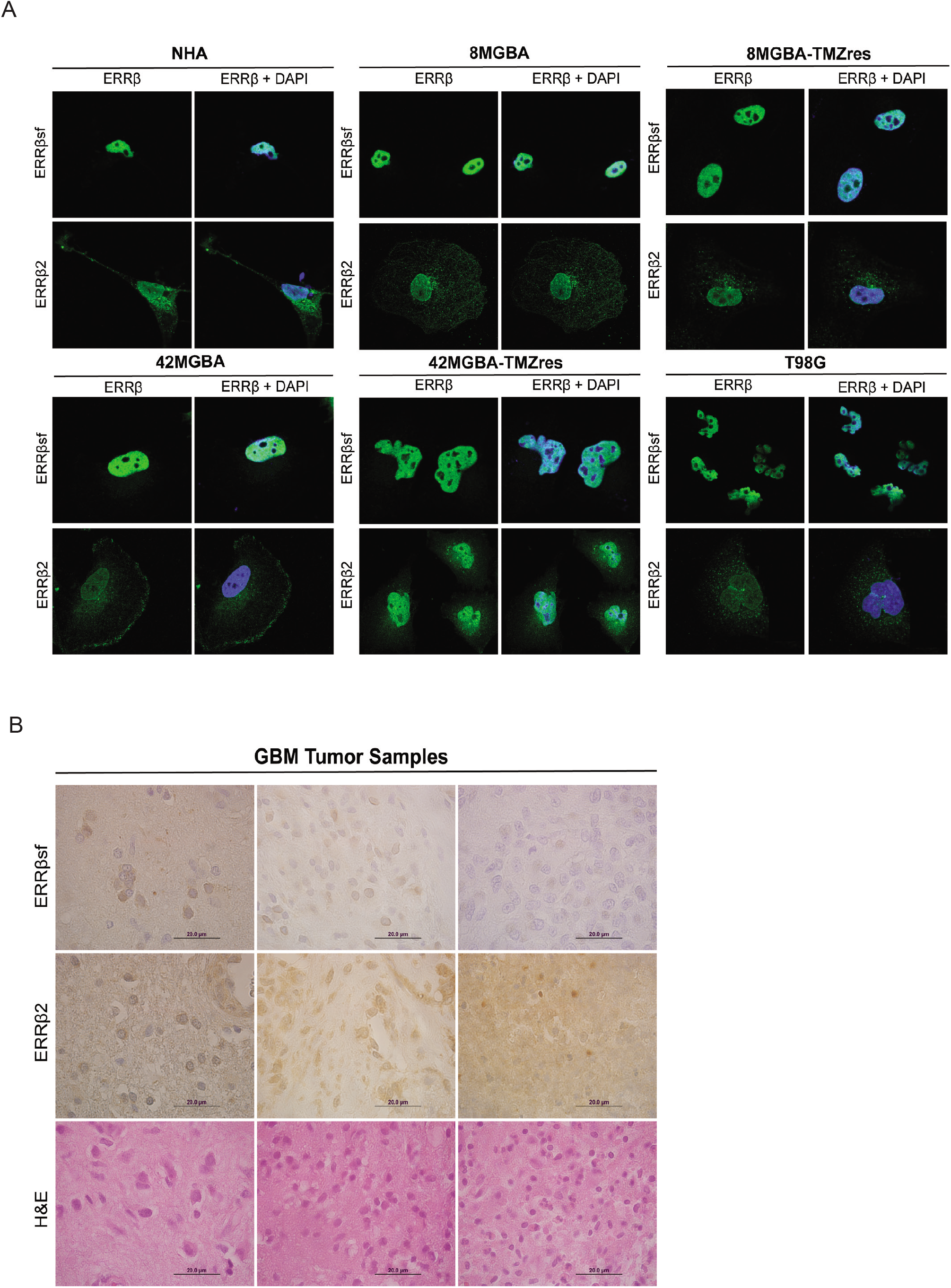
**(A)** Immunofluorescent staining of ERRβ2 and ERRβsf depicts nuclear localization of ERRβsf, and nuclear and cytoplasmic localization of ERRβ2 in NHAs and TMZ sensitive and resistant GBM cell lines. **(B)** Immunohistochemistry of ERRβ2 and ERRβsf also illustrates broad expression of these two isoforms in three different primary GBM tumor samples.

### The long ERRβ2 isoform suppresses GBM cell migration and interacts with the actin nucleation-promoting factor cortactin

Prior studies implicate ERRβsf as a transcription factor with activity at multiple DNA response elements, but suggest that ERRβ2 has little or no transcription factor activity and may partially repress βsf-mediated gene transcription (16, 18, 19). This knowledge, coupled with the localization of ERRβ2 expression adjacent to centrosomes and diffusely throughout the cytoplasm (Figure 2A) led us to test whether ERRβ2 might have additional functions in GBM cell migration. Knockdown of ERRβ2, but not ERRβsf, significantly enhances T98G cell migration in wound healing (**Figure 3A,B**) and transwell migration assays (**Supplementary Figure 2**).

**Figure 3.**
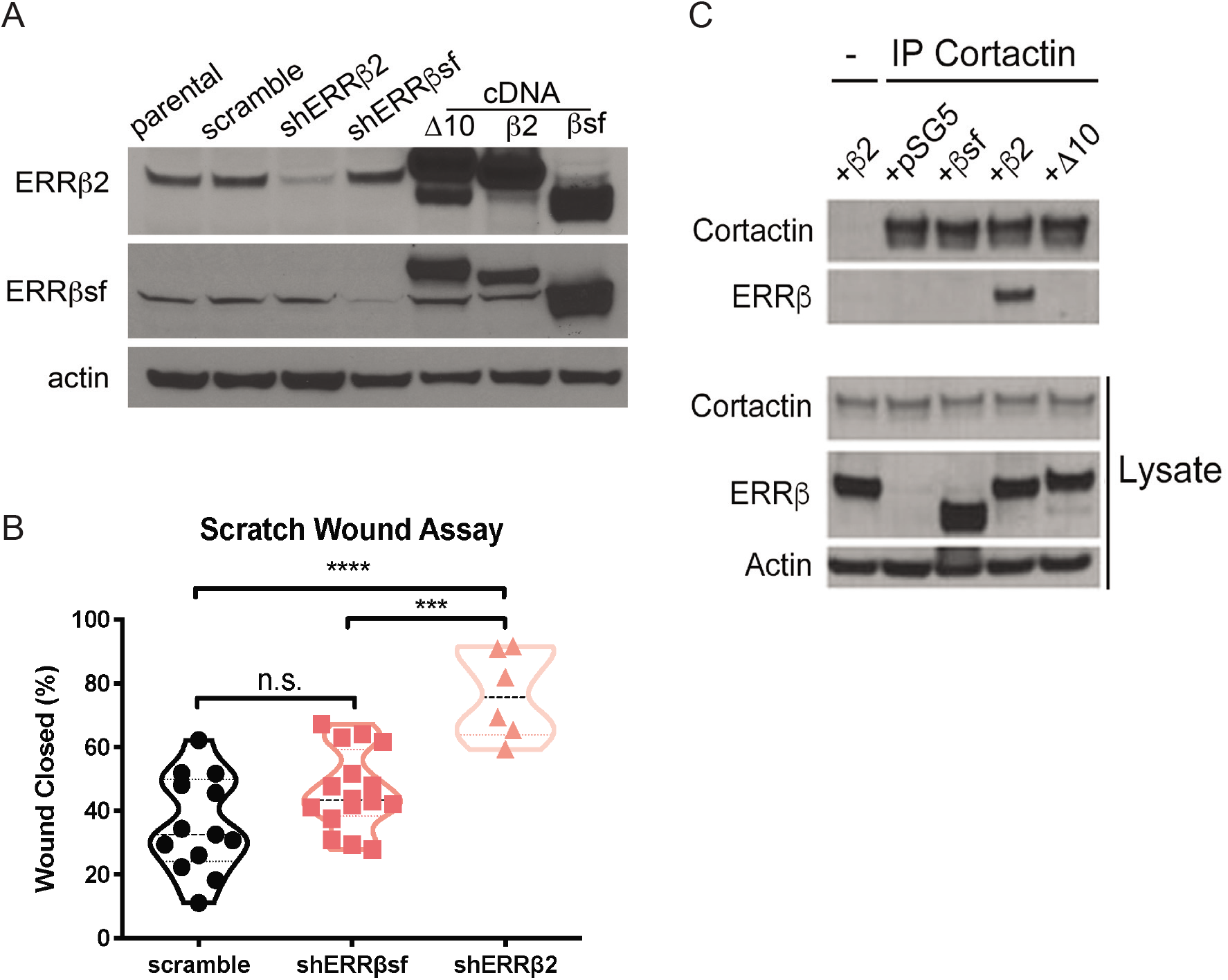
**(A, B)** Knockdown of ERRβ2, but not ERRβsf, significantly enhances T98G cell migration, as measured by scratch wound assay. In A, cells stably transduced with the indicated shRNA are compared to parental cells, or those stably transduced with a scrambled shRNA control. Lanes labeled Δ10, β2, and βsf denote lysate from cells transfected with the indicated cDNA construct. In B, data are presented as the median (line) ± minimum/maximum for 3-8 fields of view, in each of at least two independent biological replicates. Data were analyzed by one-way ANOVA with *post hoc* Tukey’s multiple comparisons test. ***, **** denote p<0.001 and p<0.0001, respectively. n.s. denotes not significant. (C) ERRβ2, but not ERRβsf or ERRβ-Δ10, forms a complex with endogenous cortactin as analyzed by immunoprecipitation. - denotes control immunoprecipitation of ERRβ2-transfected cell lysates.

F domains are carboxyl terminal extensions of the ligand binding domain of nuclear receptors, which can modify transcription factor function and recruit distinct coregulatory proteins (17). Inspection of the 67 amino acid extended carboxyl-terminal F domain unique to ERRβ2 shows a number of putative protein-protein interaction motifs and post-translational modification sites (**Supplementary Table 2**, (22)). Of these, a proline-rich region consisting of amino acids 467-472 (PLPPPP) forms the core of the consensus binding motif for the Src homology 3 (SH3) domain of the cytoskeletal protein and actin nucleation-promoting factor cortactin (35, 36). Cortactin mRNA expression is increased with increasing severity of gliomas, including GBM (**Supplementary Figure 3A**), and is significantly enriched at the leading edge of tumors (**Supplementary Figure 3B**, (26)). Cortactin has previously been implicated in GBM cell migration (37), and is expressed across all GBM cell lines (Figure 1B). Transfection of GBM cells with ERRβ cDNA shows that ERRβ2, but not ERRβsf, forms a complex with endogenous cortactin (**Figure 3C**). Together, these data suggest that the ERRβ2 isoform plays a role in cell migration and is uniquely capable of binding cortactin.

### ERRβ activation by DY131 suppresses GBM cell migration and remodels the actin cytoskeleton

We next examined the impact of activating endogenous ERRβ on actin cytoskeletal remodeling and cell migration in GBM cells. Treatment with the ERRβ agonist DY131 has no effect on ERRβ2, ERRβsf, cortactin, or MGMT protein expression, but modestly increases cleavage of PARP in 42MBGA-TMZres and T98G cell lines, indicative of apoptosis (**Figure 4A**). However, DY131 induces a marked redistribution of cortactin away from the cell membrane and towards the center of the cell body in multiple GBM cell lines (**Figure 4B**, white arrowheads vs. asterisks). In two of three TMZ resistant GBM cell lines (42MGBA-TMZres and T98G), DY131 also significantly increases F-actin polymers, as measured by fluorescence-activated cell sorting for F-actin (**Figure 4C**). Loss of cortactin membrane localization coupled with increased F-actin polymers can be indicative of impaired cell movement (38, 39), and consistent with this we show that DY131 significantly inhibits the migration of multiple GBM cell lines (**Figure 4D**).

**Figure 4.**
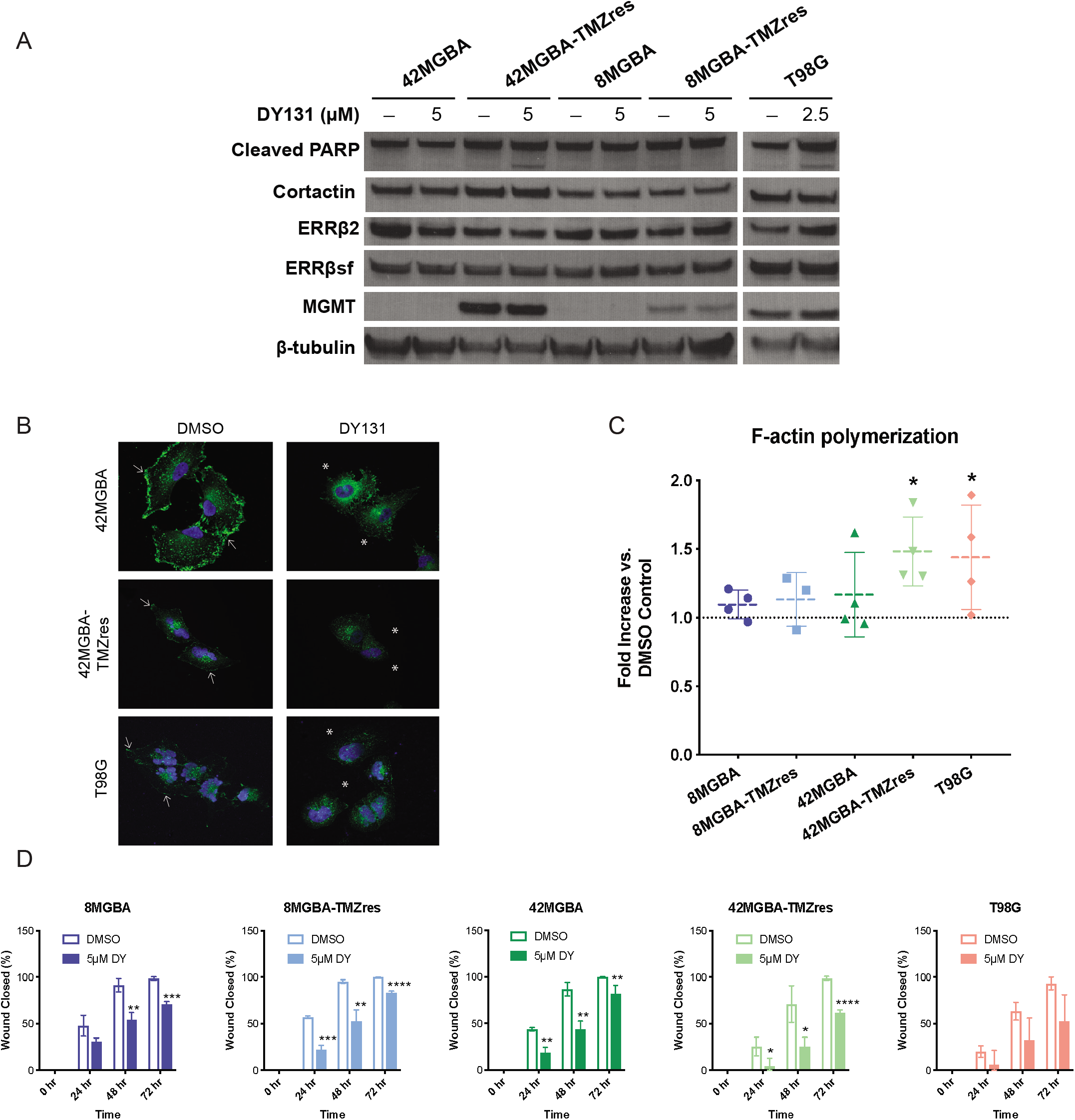
**(A)** Western blot analysis of PARP cleavage, cortactin, ERRβ2, ERRβsf, and MGMT expression in designated cell lines treated with DY131 or DMSO control (denoted as -) for 24 hours. **(B)** Immunofluorescent staining of cortactin in designated cell lines treated with 5 μM (42MBGA and 42MBGA-TMZres) or 2.5 μM DY131 (T98G) or DMSO control for 24 hours prior to fixation, staining, and imaging. DY131 treatment causes a redistribution of cortactin away from the membrane (DMSO, white arrowheads vs. DY131, white asterisks). **(C)** Changes in F-actin polymerization were measured by flow cytometry analysis of cells stained with Acti-Stain 488 phalloidin treated with 5 μM DY131 or DMSO control for 24 hours. Fold increase in F-actin polymerization in each DY131-treated cell line relative to its own DMSO control are presented as mean ± SD for 3-4 independent biological replicates. Data were analyzed by Mann-Whitney test, where * denotes p<0.05. **(D)** Scratch-wound analysis of two-dimensional migration over 72 hours of treatment with the indicated concentration of DY131 or DMSO control. Data are presented as mean ± SD for 3 independent biological replicates. Data were analyzed by Mann-Whitney test at each time point, where *, **, ***, **** denote p<0.05, p<0.01, p<0.001, and p<0.0001, respectively.

### Inhibition of Cdc2-like kinases (CLKs) shifts ERRβ isoform expression and potentiates DY131-mediated inhibition of growth and migration in GBM cells

ERRβ2 and ERRβsf have distinct roles in transcription factor activity and differentially suppress tumor cell growth and motility, but how splicing of ERRβ (gene = *ESRRB*) is regulated to produce these isoforms is unknown. The spliceosome is the primary driver of differential splicing, but heterogeneous nuclear ribonucleoprotein (hnRNP) and serine/arginine rich splicing factor (SRSF, SR protein) families cooperate with the spliceosome to enhance the exclusion vs. inclusion (respectively) of specific exons (**Figure 5A**). SR proteins broadly promote exon inclusion by binding *cis* exon splicing enhancers (ESEs), while hnRNPs are able to antagonize SR proteins, which promotes exon skipping (e.g. (40)). The ERRβ2 transcript has a unique exon inclusion event (exon 10), and multiple high-confidence SRSF6 consensus ESEs are predicted in *ESRRB* exon 10 (**Figure 5B**) (27, 28).

**Figure 5.**
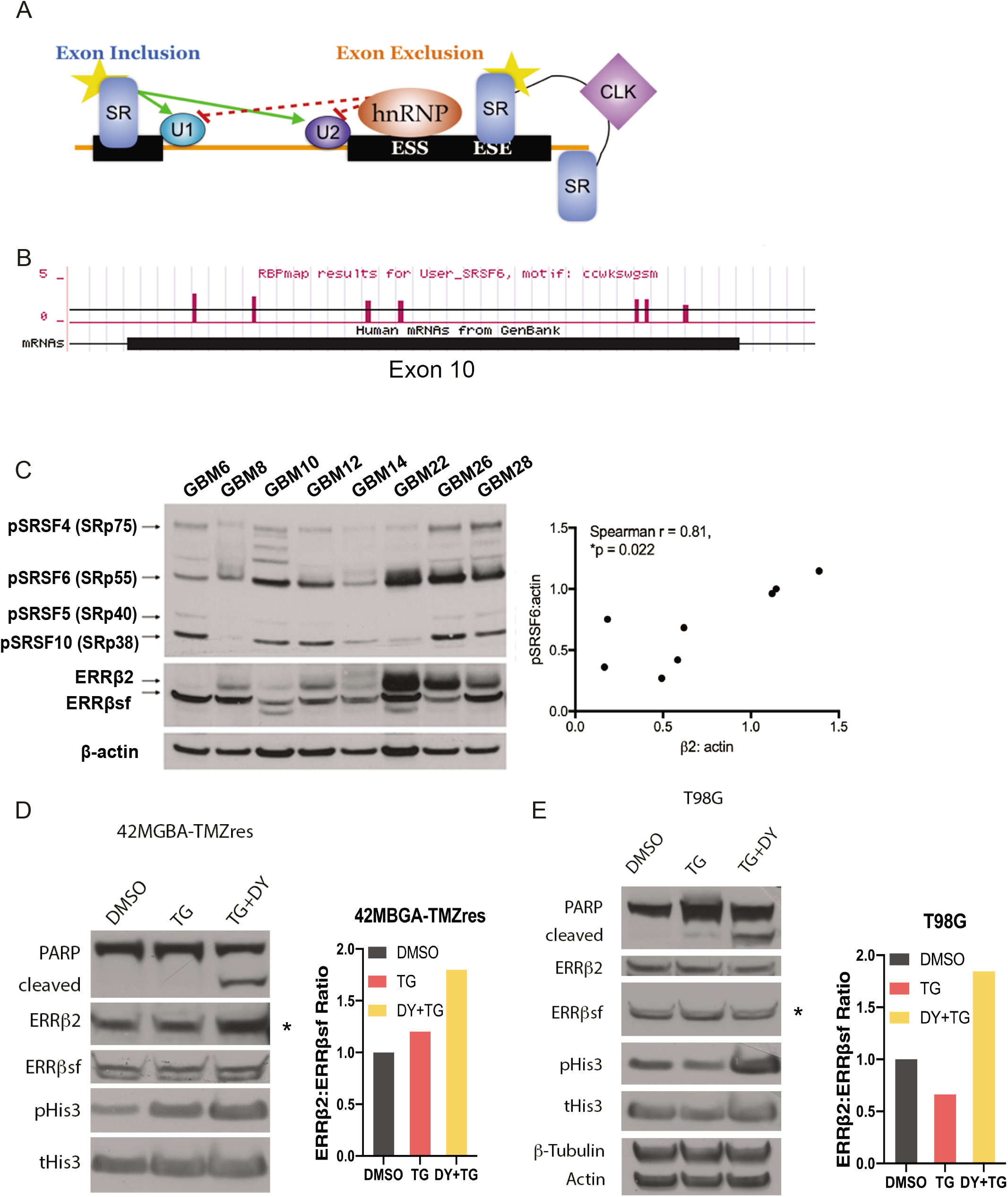
**(A)** Simplified schematic showing interplay between SR proteins and hnRNPs, which broadly promote exon inclusion or exclusion, respectively. CLK = cdc2-like kinase, star = phosphorylation. U1,2 = spliceosomal components. ESE = exon splicing enhancer, ESS = exon splicing suppressor. **(B)** RBPmap prediction of high-confidence SRSF6 binding sites in ERRβ2-specific exon 10. **(C)** Representative Western blot of pSRs (1H4), ERRβ2 and ERRβsf expression, and Spearman rank-order correlation for pSRSF6 with ERRβ2, but not ERRβsf, in GBM PDXs. Arrows indicate ERRβ2 and pSRSF6. **(D, E)** Changes in ERRβ2 and ERRβsf expression, PARP cleavage, and Serine 10 phosphorylation of histone H3 were measured in 42MGBA-TMZres (D) and T98G (E) cells treated with 50 μM TG-003, the combination of 50 μM TG-003 and 5 μM DY131, or DMSO for 24 hours. Asterisks indicate increased ERRβ2 (D) or decreased ERRβsf expression (E), both of which serve to increase the relative of abundance of ERRβ2. Bar charts depict the normalized ratio of ERRβ2 to ERRβsf protein expression by densitometry.

Serine phosphorylation (p) of SR proteins modifies their ability to promote exon inclusion. Hyper-phosphorylated SR proteins are recruited to sites of active transcription and pre-mRNA processing, where they can bind ESEs and strengthen splicing recognition sites (7). After exon-exon joining in the now-mature transcript, SR proteins become hypo-phosphorylated and are exported from the nucleus (41). Using a pan-pSR antibody, we show there is a significant positive correlation between pSRSF6 and the ERRβ2 isoform in GBM PDX models (**Figure 5C**), but no correlation between pSRSF6 and ERRβsf, or ERRβ2 and any other pSR protein. One of the kinase families responsible for catalyzing nuclear phosphorylation of SR proteins is the cdc2-like kinase (CLK) family (41). TG-003 is a pan-CLK-1, −2, and −4 inhibitor that suppresses the phosphorylation of multiple SR proteins, leading to differential splicing of a broad range of responsive mRNAs (42, 43). We therefore tested TG-003 in two TMZ-resistant GBM models as a strategy to shift the balance of ERRβ2 vs. ERRβsf isoforms (**Figure 5D, E**). In 42MGBA-TMZres cells, the combination of TG-003 and DY131 markedly upregulates ERRβ2 expression (Figure 5D, asterisk and bars), thereby increasing the ERRβ2:ERRβsf ratio. In T98G cells, the combination of TG-003 and DY131 also increases the ratio of ERRβ2:ERRβsf, although here this is achieved through decreased expression of the ERRβsf isoform (Figure 5E, asterisk and bars). Differential patterns of SR protein expression and phosphorylation are observed in these cell models (**Supplementary Figure 4**). In both cell lines, the combination of TG-003 and DY131 enhances PARP cleavage and Serine 10 phosphorylation of histone H3, which are suggestive of increased apoptosis and of mitotic arrest, respectively.

As pan-CLK inhibition shifts isoform expression in favor of ERRβ2, and ERRβ2 has a more pronounced pro-apoptotic, anti-mitotic, and anti-migratory activity ((19), and Figures 3B, 5D, 5E, and Supplementary Figure 2), we tested whether combination treatment with TG-003 and DY131 would more robustly induce G2/M arrest, apoptosis, and inhibit cell migration across all of our GBM models. Multiple TMZ sensitive and resistant GBM cell lines exhibit significantly enhanced cell death and mitotic arrest in response to combination treatment with TG-003 and DY131 (**Figure 6A, B, and Supplementary Figure 5**), with 42MGBA-TMZres and T98G cells being the most sensitive. Importantly, T98G cells treated with the combination of TG-003 and DY131 undergo equivalent G2/M arrest to T98G cells in which ERRβsf has been knocked down (**Supplementary Figure 5**), further suggesting that this phenotype is driven predominantly by ERRβ2. By contrast, MO3.13 oligodendrocytes showed no significant induction of G2/M arrest or cell death by either drug alone or the combination. In scratch-wound assays using dose-reduced concentrations of TG-003 and DY131 to isolate specific effects on migration vs. general cell viability, combination treatment significantly inhibits cell migration in multiple TMZ sensitive and resistant GBM cell lines (**Figure 6C**).

**Figure 6.**
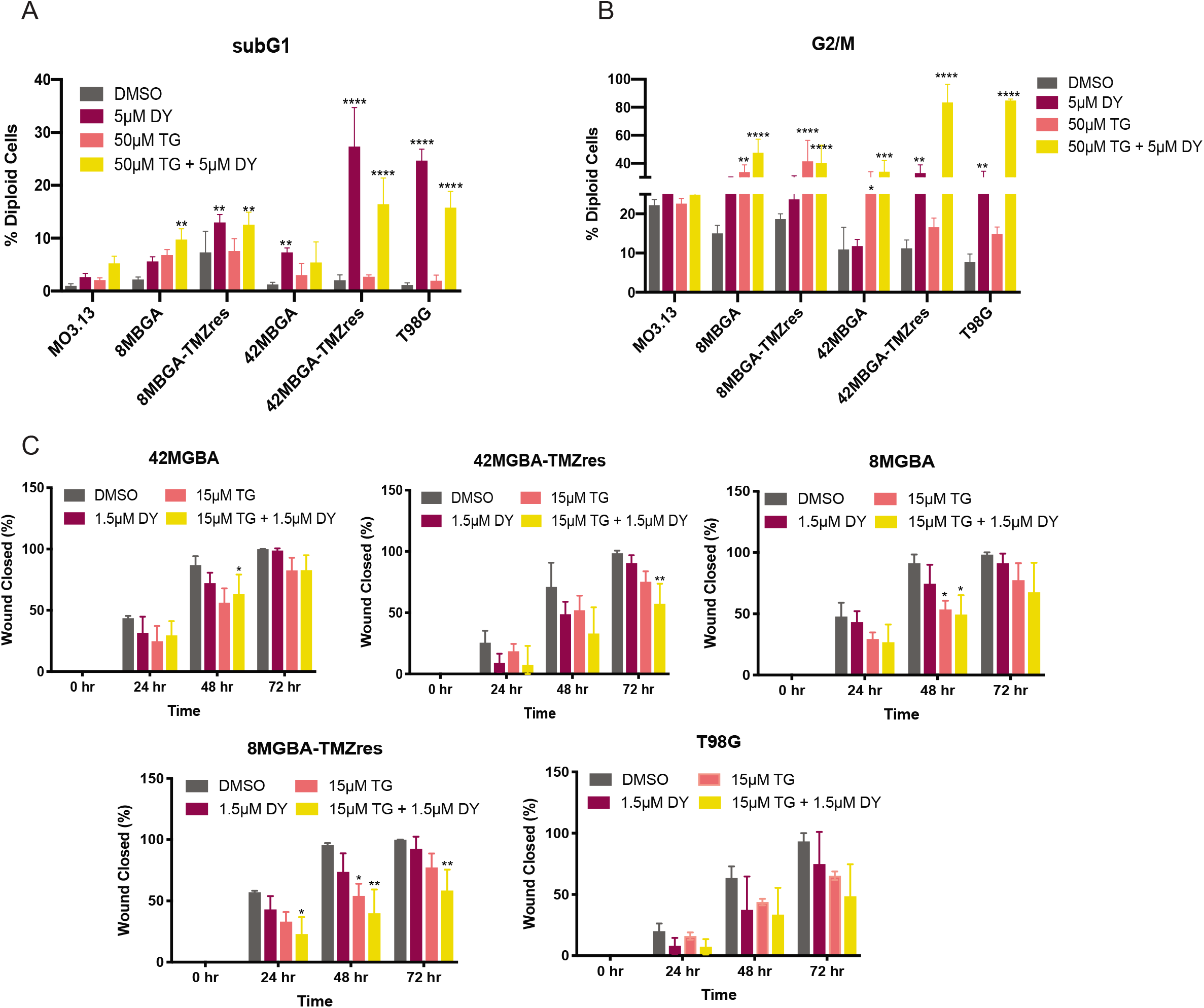
**(A, B)** Flow cytometric cell cycle analysis of SubG1 (fragmented DNA, A) and G2/M (B) fractions of immortalized oligodendrocyte MO3.13 and GBM cell lines treated with 5 μM DY131, 50 μM TG-003, the combination of 5 μM DY131 + 50 μM TG-003 (TGDY), or DMSO control for 24 hours. For each panel, data are presented as the mean ± SD for 3 independent biological replicates. Data were analyzed by one-way ANOVA with *post hoc* Tukey’s multiple comparisons test. *,**,***,**** denote p<0.05, p<0.01, p<0.001, and p<0.0001, respectively. **(C)** Scratch-wound analysis of two-dimensional migration over 72 hours of treatment with dose-reduced concentrations of DY131 and TG-003, as indicated in the legend. For each panel, data are presented as the mean ± SD for 3 independent biological replicates. Data were analyzed by one-way ANOVA at each time point with *post hoc* Tukey’s multiple comparisons test. *,**,***,**** denote p<0.05, p<0.01, p<0.001, and p<0.0001, respectively.

### Combination treatment with TG-003 and DY131 inhibits the growth of GBM intracranial xenografts

A key limitation of novel treatment strategies for GBM is failure of therapeutic agents to effectively penetrate the blood-brain barrier (BBB). The BBB protects the central nervous system from a range of endogenous and exogenous insults, and while this can be locally compromised by the primary tumor, disseminated GBM cells remaining after surgical resection are largely shielded from systemic therapies. DY131 and TG-003 are both reported to be brain penetrant in mice (9, 14, 15), so we tested these drugs individually and in combination using a zebrafish intracranial xenograft model (**Figure 7A**). The zebrafish BBB forms at ~3 days postfertilization (dpf) and is functionally similar to that of higher organisms (44, 45). We modified published intracranial xenograft procedures (**Figure 7B**, (46, 47)), injecting labeled 42MGBA-TMZres cells into 1.5 dpf zebrafish embryos. Two to 3 days after the BBB forms (by 6 dpf), tumors were imaged, fish were treated daily by addition of test compounds to the fish water, and tumors were imaged again at 10 dpf. Comparing the area of pre-vs. post-treatment confocal tumor images shows that 54% of tumors in control-treated fish increased in size, but 5 days of treatment with the dose-reduced combination of TG-003 (15 μM) and DY131 (1.5 μM) reduces this to 32% (**Figure 7C**, Cochran-Armitage p=0.09). In parallel, Cascade Blue-conjugated dextran imaging at 10 dpf was used to assess the integrity of the BBB. In both DMSO control- and combination-treated fish, signal is occluded from the brain (**Figure 7D**). Collectively, these results support penetrance of TG-003 and DY131 with an intact BBB, and provide evidence of anti-tumor activity of this combination *in vivo* against TMZ-resistant GBM.

**Figure 7.**
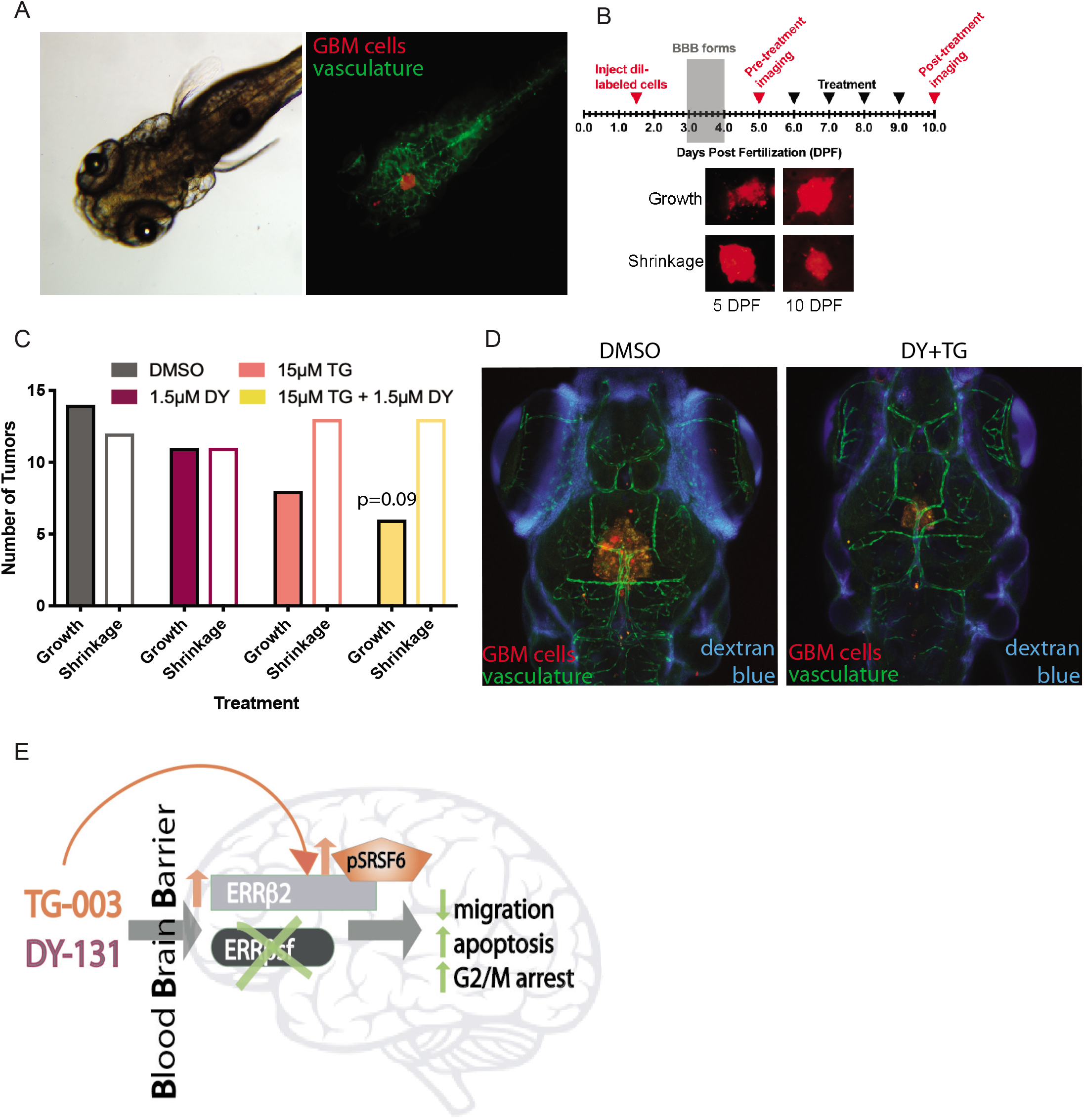
**(A)** Brightfield (left) and fluorescent (right) images of 5 day post-fertilization (dpf) zebrafish bearing xenografts of 42MGBA-TMZres cells. DiI-labeled tumor cells are red and the vasculature is green. **(B)** Experimental schematic for *in vivo* zebrafish model with representative images of DiI-labeled 42MBGA-TMZres cells pre- and post-treatment. **(C)** Growth or shrinkage of 42MGBA-TMZres xenografts following 5 days of treatment with 1.5 μM DY131, 15 μM TG-003, the combination of 1.5 μM DY131 + 15 μM TG-003, or DMSO shows a trend towards tumor shrinkage in the combined treatment group. Data are presented as fraction of fish showing tumor growth vs. tumor shrinkage for 19-26 fish per group. Data were analyzed by χ^2^ test for trend (Cochran-Armitage), p=0.09. **(D)** Fluorescent images of 10 day post-fertilization (dpf) zebrafish bearing xenografts of 42MGBA-TMZres cells following 5 days of treatment with DMSO control (left) or the combination of 1.5 μM DY131 + 15 μM TG-003 (right). DiI-labeled tumor cells are red, the vasculature is green, and Cascade Blue-conjugated dextran shows integrity of the blood-brain barrier (BBB). **(E)** Summary schematic of key findings. CLK inhibition shifts isoform expression in favor of ERRβ2, and potentiates ERRβ agonist-mediated inhibition of growth and migration in GBM.

## DISCUSSION

Here, we show that the ERRβ agonist, DY131, is growth inhibitory in a broad set of GBM cell lines. The cytoplasmic and nuclear ERRβ2 isoform is expressed in multiple GBM cell lines, PDX models, and human tumors, where it suppresses GBM cell migration, and interacts with the actin nucleation-promoting factor cortactin. Treatment with the ERRβ agonist DY131 remodels the actin cytoskeleton and suppresses migration, and we further show that broad CLK inhibition shifts isoform expression in favor of ERRβ2, and potentiates ERRβ agonist-mediated inhibition of growth and migration in GBM cells. Combined treatment with DY131 and TG-003 also produces a non-significant trend toward inhibition of intracranial tumor growth.

ERRβ2, unlike ERRβsf, is specifically expressed in primates and localized to both the cytoplasm and the nucleus. Altered subcellular localization of splice variants is a common phenotype, especially in brain cancer and brain development (e.g. (48)), and suggests a specialized function of the cytoplasmic primate-specific ERRβ2 nuclear receptor isoform. Inclusion of a unique F domain that may allow ERRβ2 to bind cortactin, an essential regulator of the actin cytoskeleton and cell motility in GBM, gives further insight into its specific function. GBM is a highly migratory tumor in which cortactin plays a critical role (37). We show that the ERRβ agonist DY131 decreases migration, while shRNA-mediated silencing of ERRβ2, but not ERRβsf, increases the migration of GBM cells in the absence of this agonist. This suggests that changing the balance between ERRβ2 and ERRβsf could successfully limit GBM migration and local invasion. However, si/shRNA-dependent strategies have many limitations and few clinical successes (49). Splice-switching antisense oligonucleotides (AONs) designed to modify isoform expression are emerging as useful approaches in musculoskeletal disorders (50), but to our knowledge they have not yet been tested in GBM.

In the absence of established nucleotide-based strategies, we looked to splicing modulatory drugs, specifically splicing kinase inhibitors, to increase the ERRβ2:ERRβsf ratio. We show that the pan-CLK inhibitor TG-003 in combination with DY131 increases the relative expression of the ERRβ2 isoform in two TMZ-resistant GBM cells, and causes a more robust inhibition of G2/M arrest and migration which have previously been shown to be ERRβ2-dependent. Coupled with the knowledge that differential splicing is a key driver of phenotypic diversity and plasticity in the malignant brain (4–6), our results suggest a new paradigm in which splicing modulatory drugs could be an effective approach to inhibit GBM migration and invasion by changing the balance of pro-vs. anti-migratory isoforms of ERRβ and other proteins. TG-003 is a broad spectrum CLK inhibitor, and we do not yet know which of the CLKs are most important for shifting isoform expression in favor of ERRβ2. Sakuma *et al*. showed that the strength of uracil-rich polypyrimidine tracts, exon length, and abundance of splicing factor binding sites characterize TG-003-responsive exons in human and mouse skeletal muscle cells (51). Additional studies are necessary to determine how ESRRB sequence features and the repertoire of RNA binding factors beyond SR proteins dictate context-dependent pre-mRNA processing and the expression of distinct ERRβ isoforms. Nevertheless, we provide evidence that ERRβ, specifically the ERRβ2 isoform, has pro-apoptotic and anti-migratory functions in GBM, and that splicing modulatory drugs such as CLK inhibitors are a novel strategy for shifting the balance of ERRβ isoforms to potentiate ERRβ agonist-mediated inhibition of growth and migration in GBM cells and intracranial tumors (**Figure 7E**).

Poor brain penetrance is also a common cause of failure for novel therapeutic agents in GBM clinical studies despite apparently robust anti-tumor activity *in vitro* (3), which is why intracranial GBM models are an essential preclinical test of candidate small molecules and treatment strategies. We show that the combination of DY131 and TG-003 numerically (but not significantly) restricts the growth of TMZ-resistant intracranial GBM xenografts in a context in which the the BBB remains intact. Zebrafish are an ideal model organism for early-phase screening of GBM intracranial xenografts, due to their evolutionarily conserved BBB structure, capacity for intrinsic neovascularization of implanted tumors, and in pigmentation-deficient transgenic animals, tumor development and response to treatment can be tracked in real time in live animals (52). While others (9, 14, 15) show that DY131 and TG-003 are brain penetrant and our data demonstrate that these small molecules may have anti-tumor activity in combination, further optimization and drug development efforts will be necessary to improve the efficacy of these and other splicing modulatory drugs and nuclear receptor ligands.

A potential limitation of our study is that, aside from co-immunoprecipitation studies in Figure 3C, it does not address the contribution of ERRβ-Δ10 to GBM. The field currently lacks reagents that are selective enough to study the endogenous function of the ERRβ-Δ10 isoform. The extended carboxyl-terminal F domain of the exon 10-deleted (ERRβ-Δ10) isoform has a proline-rich sequence that conforms more closely to the consensus binding motif for the SH3 domain-containing protein amphiphysin (Supplementary Table 2) (53), yet we show that exogenously expressed ERRβ-Δ10 is unable to interact with endogenous cortactin. Previously published studies in cells transfected with cDNA for ERRβ-Δ10 show that this isoform, like ERRβsf, localizes to the nucleus (and not the cytoplasm), and has transcription factor activity (16). A second limitation is that ERRβ agonist ligands can have activity against other members of the estrogen-related receptor family, or off-target effects. DY131 is also an agonist for the closely related ERRγ (12, 54), and a prior report implicates this compound as an antagonist of Hedgehog signaling (55). However, we have shown previously that shRNA-mediated silencing of ERRγ does not abrogate DY131-mediated phenotypes, and the Hedgehog pathway inhibitors cyclopamine and vismodegib do not phenocopy DY131 in GBM cells (19). A third limitation of the study is the non-significant trend toward inhibition of intracranial tumor growth by the DY131 + TG-003 combination. One potential explanation is that zebrafish viability is optimal at 28°C and human GBM cell lines are adapted to grow at 37°C, so a typical compromise employed by zebrafish xenograft studies (including ours) is to maintain the fish at 33°C. This may contribute to both poor tolerance of the embryos to intracranial injection and the shrinkage shown in the DMSO treated cells. Orthotopic injections in immunocompromised mouse models are an alternate approach and future strategy that will be considered to circumvent this limitation.

In summary, our current work builds upon prior studies that first established the more robust anti-tumor effects of the ERRβ2 isoform. We demonstrate that ERRβ2 is broadly expressed in multiple GBM cell lines and PDXs, plays an anti-migratory role that we hypothesize is mediated by its interaction with cortactin via a unique F domain, and induces a robust G2/M arrest and apoptosis upon treatment with the ERRβ agonist DY131. We further demonstrate the efficacy of splicing modulatory drugs to shift the ERRβ2:ERRβsf isoform ratio in lieu of established AON-dependent approaches, or sh/siRNA-mediated isoform targeting that has thus far been unsuccessfully clinically in cancer. Lastly, we show preliminary potential for combination treatment with DY131 and TG-003 to shrink tumor growth *in vivo*.

## Conflict of Interest and Author Contribution Statement

The authors declare no potential conflict of interest. DMT contributed to study design, performed experiments, analyzed data, and wrote the paper. SAK performed experiments, analyzed data, and wrote the paper. CJT performed experiments and analyzed data. MMH performed experiments. SDD performed experiments. JNS provided patient-derived xenograft samples. EG contributed to study design, and performed experiments. RBR contributed to study design, performed experiments, analyzed data, and wrote the paper. All authors reviewed, edited, and approved the manuscript.

## ACKNOWLEDGEMENTS

We wish to thank members of the Riggins laboratory as well as Drs. Maria Laura Avantaggiati, Deborah Berry, Karen Creswell, Brent Harris, Peter Johnson, Supti Sen, Alexandra Taraboletti, Jeffrey Toretsky, Todd Waldman, and Dan Xun for sharing reagents, scientific insights, technical assistance, and/or editorial comments on the manuscript.

## REFERENCES

1. Perazzoli, G., Prados, J., Ortiz, R., Caba, O., Cabeza, L., Berdasco, M., Gónzalez, B., and Melguizo, C. (2015) Temozolomide Resistance in Glioblastoma Cell Lines: Implication of MGMT, MMR, P-Glycoprotein and CD133 Expression. PLoS One 10, e0140131

2. Prados, M. D., Byron, S. A., Tran, N. L., Phillips, J. J., Molinaro, A. M., Ligon, K. L., Wen, P. Y., Kuhn, J. G., Mellinghoff, I. K., de Groot, J. F., Colman, H., Cloughesy, T. F., Chang, S. M., Ryken, T. C., Tembe, W. D., Kiefer, J. A., Berens, M. E., Craig, D. W., Carpten, J. D., and Trent, J. M. (2015) Toward precision medicine in glioblastoma: the promise and the challenges. Neuro Oncol

3. Parrish, K. E., Sarkaria, J. N., and Elmquist, W. F. (2015) Improving drug delivery to primary and metastatic brain tumors: strategies to overcome the blood-brain barrier. Clin Pharmacol Ther 97, 336–346

4. Braun, C. J., Stanciu, M., Boutz, P. L., Patterson, J. C., Calligaris, D., Higuchi, F., Neupane, R., Fenoglio, S., Cahill, D. P., Wakimoto, H., Agar, N. Y. R., Yaffe, M. B., Sharp, P. A., Hemann, M. T., and Lees, J. A. (2017) Coordinated Splicing of Regulatory Detained Introns within Oncogenic Transcripts Creates an Exploitable Vulnerability in Malignant Glioma. Cancer Cell 32, 411–426 e411

5. Golan-Gerstl, R., Cohen, M., Shilo, A., Suh, S. S., Bakàcs, A., Coppola, L., and Karni, R. (2011) Splicing factor hnRNP A2/B1 regulates tumor suppressor gene splicing and is an oncogenic driver in glioblastoma. Cancer Res 71, 4464–4472

6. Ferrarese, R., Harsh, G. R., Yadav, A. K., Bug, E., Maticzka, D., Reichardt, W., Dombrowski, S. M., Miller, T. E., Masilamani, A. P., Dai, F., Kim, H., Hadler, M., Scholtens, D. M., Yu, I. L., Beck, J., Srinivasasainagendra, V., Costa, F., Baxan, N., Pfeifer, D., von Elverfeldt, D., Backofen, R., Weyerbrock, A., Duarte, C. W., He, X., Prinz, M., Chandler, J. P., Vogel, H., Chakravarti, A., Rich, J. N., Carro, M. S., and Bredel, M. (2014) Lineage-specific splicing of a brain-enriched alternative exon promotes glioblastoma progression. J Clin Invest 124, 2861–2876

7. Shepard, P. J., and Hertel, K. J. (2009) The SR protein family. Genome Biol 10, 242

8. Park, S. Y., Piao, Y., Thomas, C., Fuller, G. N., and de Groot, J. F. (2016) Cdc2-like kinase 2 is a key regulator of the cell cycle via FOXO3a/p27 in glioblastoma. Oncotarget 7, 26793–26805

9. Bidinosti, M., Botta, P., Krüttner, S., Proenca, C. C., Stoehr, N., Bernhard, M., Fruh, I., Mueller, M., Bonenfant, D., Voshol, H., Carbone, W., Neal, S. J., McTighe, S. M., Roma, G., Dolmetsch, R. E., Porter, J. A., Caroni, P., Bouwmeester, T., Lüthi, A., and Galimberti, I. (2016) CLK2 inhibition ameliorates autistic features associated with SHANK3 deficiency. Science 351, 1199–1203

10. Seiler, M., Yoshimi, A., Darman, R., Chan, B., Keaney, G., Thomas, M., Agrawal, A. A., Caleb, B., Csibi, A., Sean, E., Fekkes, P., Karr, C., Klimek, V., Lai, G., Lee, L., Kumar, P., Lee, S. C., Liu, X., Mackenzie, C., Meeske, C., Mizui, Y., Padron, E., Park, E., Pazolli, E., Peng, S., Prajapati, S., Taylor, J., Teng, T., Wang, J., Warmuth, M., Yao, H., Yu, L., Zhu, P., Abdel-Wahab, O., Smith, P. G., and Buonamici, S. (2018) H3B-8800, an orally available small-molecule splicing modulator, induces lethality in spliceosome-mutant cancers. Nat Med 24, 497–504

11. Giguère, V., Yang, N., Segui, P., and Evans, R. M. (1988) Identification of a new class of steroid hormone receptors. Nature 331, 91–94

12. Yu, D. D., and Forman, B. M. (2005) Identification of an agonist ligand for estrogen-related receptors ERRbeta/gamma 1. Bioorg.Med.Chem.Lett. 15, 1311–1313

13. Zuercher, W. J., Gaillard, S., Orband-Miller, L. A., Chao, E. Y., Shearer, B. G., Jones, D. G., Miller, A. B., Collins, J. L., McDonnell, D. P., and Willson, T. M. (2005) Identification and structure-activity relationship of phenolic acyl hydrazones as selective agonists for the estrogen-related orphan nuclear receptors ERRbeta and ERRgamma 2 7337. J.Med.Chem. 48, 3107–3109

14. Byerly, M. S., Al Salayta, M., Swanson, R. D., Kwon, K., Peterson, J. M., Wei, Z., Aja, S., Moran, T. H., Blackshaw, S., and Wong, G. W. (2013) Estrogen-related receptor β deletion modulates whole-body energy balance via estrogen-related receptor γ and attenuates neuropeptide Y gene expression. Eur J Neurosci

15. Byerly, M. S., Swanson, R. D., Wong, G. W., and Blackshaw, S. (2013) Estrogen-related receptor β deficiency alters body composition and response to restraint stress. BMC Physiol 13, 10

16. Zhou, W., Liu, Z., Wu, J., Liu, J. H., Hyder, S. M., Antoniou, E., and Lubahn, D. B. (2006) Identification and characterization of two novel splicing isoforms of human estrogen-related receptor beta. J Clin Endocrinol Metab 91, 569–579

17. Patel, S. R., and Skafar, D. F. (2015) Modulation of nuclear receptor activity by the F domain. Mol Cell Endocrinol 418 Pt 3, 298–305

18. Heckler, M. M., Zeleke, T. Z., Divekar, S. D., Fernandez, A. I., Tiek, D. M., Woodrick, J., Farzanegan, A., Roy, R., Üren, A., Mueller, S. C., and Riggins, R. B. (2016) Antimitotic activity of DY131 and the estrogen-related receptor beta 2 (ERRβ2) splice variant in breast cancer. Oncotarget 7, 47201–47220

19. Heckler, M. M., and Riggins, R. B. (2015) ERRβ splice variants differentially regulate cell cycle progression. Cell Cycle 14, 31–45

20. Tiek, D. M., Rone, J. D., Graham, G. T., Pannkuk, E. L., Haddad, B. R., and Riggins, R. B. (2018) Alterations in Cell Motility, Proliferation, and Metabolism in Novel Models of Acquired Temozolomide Resistant Glioblastoma. Sci Rep 8, 7222

21. Rasband, W. S. (2004) ImageJ. National Institutes of Health, Bethesda MD, http://rsb.info.nih.gov/ij/

22. Obenauer, J. C., Cantley, L. C., and Yaffe, M. B. (2003) Scansite 2.0: Proteome-wide prediction of cell signaling interactions using short sequence motifs. Nucleic Acids Res 31, 3635–3641

23. Lindsay, S. J., Xu, Y., Lisgo, S. N., Harkin, L. F., Copp, A. J., Gerrelli, D., Clowry, G. J., Talbot, A., Keogh, M. J., Coxhead, J., Santibanez-Koref, M., and Chinnery, P. F. (2016) HDBR Expression: A Unique Resource for Global and Individual Gene Expression Studies during Early Human Brain Development. Front Neuroanat 10, 86

24. Papatheodorou, I., Fonseca, N. A., Keays, M., Tang, Y. A., Barrera, E., Bazant, W., Burke, M., Fullgrabe, A., Fuentes, A. M., George, N., Huerta, L., Koskinen, S., Mohammed, S., Geniza, M., Preece, J., Jaiswal, P., Jarnuczak, A. F., Huber, W., Stegle, O., Vizcaino, J. A., Brazma, A., and Petryszak, R. (2018) Expression Atlas: gene and protein expression across multiple studies and organisms. Nucleic Acids Res 46, D246–D251

25. Goldman, M., Craft, B., Hastie, M., Repecka, K., Kamath, A., McDade, F., Rogers, D., Brooks, A. N., Zhu, J., and Haussler, D. (2019) The UCSC Xena platform for public and private cancer genomics data visualization and interpretation. bioRxiv, 326470

26. Puchalski, R. B., Shah, N., Miller, J., Dalley, R., Nomura, S. R., Yoon, J. G., Smith, K. A., Lankerovich, M., Bertagnolli, D., Bickley, K., Boe, A. F., Brouner, K., Butler, S., Caldejon, S., Chapin, M., Datta, S., Dee, N., Desta, T., Dolbeare, T., Dotson, N., Ebbert, A., Feng, D., Feng, X., Fisher, M., Gee, G., Goldy, J., Gourley, L., Gregor, B. W., Gu, G., Hejazinia, N., Hohmann, J., Hothi, P., Howard, R., Joines, K., Kriedberg, A., Kuan, L., Lau, C., Lee, F., Lee, H., Lemon, T., Long, F., Mastan, N., Mott, E., Murthy, C., Ngo, K., Olson, E., Reding, M., Riley, Z., Rosen, D., Sandman, D., Shapovalova, N., Slaughterbeck, C. R., Sodt, A., Stockdale, G., Szafer, A., Wakeman, W., Wohnoutka, P. E., White, S. J., Marsh, D., Rostomily, R. C., Ng, L., Dang, C., Jones, A., Keogh, B., Gittleman, H. R., Barnholtz-Sloan, J. S., Cimino, P. J., Uppin, M. S., Keene, C. D., Farrokhi, F. R., Lathia, J. D., Berens, M. E., Iavarone, A., Bernard, A., Lein, E., Phillips, J. W., Rostad, S. W., Cobbs, C., Hawrylycz, M. J., and Foltz, G. D. (2018) An anatomic transcriptional atlas of human glioblastoma. Science 360, 660–663

27. Jensen, M. A., Wilkinson, J. E., and Krainer, A. R. (2014) Splicing factor SRSF6 promotes hyperplasia of sensitized skin. Nat Struct Mol Biol 21, 189–197

28. Paz, I., Kosti, I., Ares, M., Cline, M., and Mandel-Gutfreund, Y. (2014) RBPmap: a web server for mapping binding sites of RNA-binding proteins. Nucleic Acids Res 42, W361–367

29. Robins, H. I., Won, M., Seiferheld, W. F., Schultz, C. J., Choucair, A. K., Brachman, D. G., Demas, W. F., and Mehta, M. P. (2006) Phase 2 trial of radiation plus high-dose tamoxifen for glioblastoma multiforme: RTOG protocol BR-0021. Neuro Oncol 8, 47–52

30. Yu, S., Wong, Y. C., Wang, X. H., Ling, M. T., Ng, C. F., Chen, S., and Chan, F. L. (2008) Orphan nuclear receptor estrogen-related receptor-beta suppresses in vitro and in vivo growth of prostate cancer cells via p21(WAF1/CIP1) induction and as a potential therapeutic target in prostate cancer. Oncogene 27, 3313–3328

31. Carlson, B. L., Pokorny, J. L., Schroeder, M. A., and Sarkaria, J. N. (2011) Establishment, maintenance and in vitro and in vivo applications of primary human glioblastoma multiforme (GBM) xenograft models for translational biology studies and drug discovery. Curr Protoc Pharmacol Chapter 14, Unit 14.16

32. Brennan, C. W., Verhaak, R. G., McKenna, A., Campos, B., Noushmehr, H., Salama, S. R., Zheng, S., Chakravarty, D., Sanborn, J. Z., Berman, S. H., Beroukhim, R., Bernard, B., Wu, C. J., Genovese, G., Shmulevich, I., Barnholtz-Sloan, J., Zou, L., Vegesna, R., Shukla, S. A., Ciriello, G., Yung, W. K., Zhang, W., Sougnez, C., Mikkelsen, T., Aldape, K., Bigner, D. D., Van Meir, E. G., Prados, M., Sloan, A., Black, K. L., Eschbacher, J., Finocchiaro, G., Friedman, W., Andrews, D. W., Guha, A., Iacocca, M., O’Neill, B. P., Foltz, G., Myers, J., Weisenberger, D. J., Penny, R., Kucherlapati, R., Perou, C. M., Hayes, D. N., Gibbs, R., Marra, M., Mills, G. B., Lander, E., Spellman, P., Wilson, R., Sander, C., Weinstein, J., Meyerson, M., Gabriel, S., Laird, P. W., Haussler, D., Getz, G., Chin, L., and Network, T. R. (2013) The somatic genomic landscape of glioblastoma. Cell 155, 462–477

33. Murnyak, B., Kouhsari, M. C., Hershkovitch, R., Kalman, B., Marko-Varga, G., Klekner, A., and Hortobagyi, T. (2017) PARP1 expression and its correlation with survival is tumour molecular subtype dependent in glioblastoma. Oncotarget 8, 46348–46362

34. Ernst, A., Hofmann, S., Ahmadi, R., Becker, N., Korshunov, A., Engel, F., Hartmann, C., Felsberg, J., Sabel, M., Peterziel, H., Durchdewald, M., Hess, J., Barbus, S., Campos, B., Starzinski-Powitz, A., Unterberg, A., Reifenberger, G., Lichter, P., Herold-Mende, C., and Radlwimmer, B. (2009) Genomic and expression profiling of glioblastoma stem cell-like spheroid cultures identifies novel tumor-relevant genes associated with survival. Clin Cancer Res 15, 6541–6550

35. Sparks, A. B., Rider, J. E., Hoffman, N. G., Fowlkes, D. M., Quillam, L. A., and Kay, B. K. (1996) Distinct ligand preferences of Src homology 3 domains from Src, Yes, Abl, Cortactin, p53bp2, PLCgamma, Crk, and Grb2. Proc Natl Acad Sci U S A 93, 1540–1544

36. MacGrath, S. M., and Koleske, A. J. (2012) Cortactin in cell migration and cancer at a glance. J Cell Sci 125, 1621–1626

37. Fujimura, A., Michiue, H., Cheng, Y., Uneda, A., Tani, Y., Nishiki, T., Ichikawa, T., Wei, F. Y., Tomizawa, K., and Matsui, H. (2013) Cyclin G2 promotes hypoxia-driven local invasion of glioblastoma by orchestrating cytoskeletal dynamics. Neoplasia 15, 1272–1281

38. Kirkbride, K. C., Sung, B. H., Sinha, S., and Weaver, A. M. (2011) Cortactin: a multifunctional regulator of cellular invasiveness. Cell Adh Migr 5, 187–198

39. Tojkander, S., Gateva, G., and Lappalainen, P. (2012) Actin stress fibers--assembly, dynamics and biological roles. J Cell Sci 125, 1855–1864

40. Rahman, M. A., Azuma, Y., Nasrin, F., Takeda, J., Nazim, M., Bin Ahsan, K., Masuda, A., Engel, A. G., and Ohno, K. (2015) SRSF1 and hnRNP H antagonistically regulate splicing of COLQ exon 16 in a congenital myasthenic syndrome. Sci Rep 5, 13208

41. Corkery, D. P., Holly, A. C., Lahsaee, S., and Dellaire, G. (2015) Connecting the speckles: Splicing kinases and their role in tumorigenesis and treatment response. Nucleus 6, 279–288

42. Muraki, M., Ohkawara, B., Hosoya, T., Onogi, H., Koizumi, J., Koizumi, T., Sumi, K., Yomoda, J., Murray, M. V., Kimura, H., Furuichi, K., Shibuya, H., Krainer, A. R., Suzuki, M., and Hagiwara, M. (2004) Manipulation of alternative splicing by a newly developed inhibitor of Clks. J Biol Chem 279, 24246–24254

43. Marcel, V., Fernandes, K., Terrier, O., Lane, D. P., and Bourdon, J. C. (2014) Modulation of p53β and p53γ expression by regulating the alternative splicing of TP53 gene modifies cellular response. Cell Death Differ

44. Jeong, J. Y., Kwon, H. B., Ahn, J. C., Kang, D., Kwon, S. H., Park, J. A., and Kim, K. W. (2008) Functional and developmental analysis of the blood-brain barrier in zebrafish. Brain Res Bull 75, 619–628

45. O’Brown, N. M., Pfau, S. J., and Gu, C. (2018) Bridging barriers: a comparative look at the blood-brain barrier across organisms. Genes Dev 32, 466–478

46. Welker, A. M., Jaros, B. D., Puduvalli, V. K., Imitola, J., Kaur, B., and Beattie, C. E. (2016) Standardized orthotopic xenografts in zebrafish reveal glioma cell-line-specific characteristics and tumor cell heterogeneity. Dis Model Mech 9, 199–210

47. Welker, A. M., Jaros, B. D., An, M., and Beattie, C. E. (2017) Changes in tumor cell heterogeneity after chemotherapy treatment in a xenograft model of glioblastoma. Neuroscience 356, 35–43

48. Lee, J. A., Damianov, A., Lin, C. H., Fontes, M., Parikshak, N. N., Anderson, E. S., Geschwind, D. H., Black, D. L., and Martin, K. C. (2016) Cytoplasmic Rbfox1 Regulates the Expression of Synaptic and Autism-Related Genes. Neuron 89, 113–128

49. Das, M., Musetti, S., and Huang, L. (2018) RNA Interference-Based Cancer Drugs: The Roadblocks, and the “Delivery” of the Promise. Nucleic Acid Ther

50. Lim, K. R. Q., and Yokota, T. (2018) Invention and Early History of Exon Skipping and Splice Modulation. Methods Mol Biol 1828, 3–30

51. Sakuma, M., Iida, K., and Hagiwara, M. (2015) Deciphering targeting rules of splicing modulator compounds: case of TG003. BMC Mol Biol 16, 16

52. Barriuso, J., Nagaraju, R., and Hurlstone, A. (2015) Zebrafish: a new companion for translational research in oncology. Clin Cancer Res 21, 969–975

53. Grabs, D., Slepnev, V. I., Songyang, Z., David, C., Lynch, M., Cantley, L. C., and De Camilli, P. (1997) The SH3 domain of amphiphysin binds the proline-rich domain of dynamin at a single site that defines a new SH3 binding consensus sequence. J Biol Chem 272, 13419–13425

54. Yu, S., Wang, X., Ng, C. F., Chen, S., and Chan, F. L. (2007) ERRgamma suppresses cell proliferation and tumor growth of androgen-sensitive and androgen-insensitive prostate cancer cells and its implication as a therapeutic target for prostate cancer. Cancer Res 67, 4904–4914

55. Wang, Y., Arvanites, A. C., Davidow, L., Blanchard, J., Lam, K., Yoo, J. W., Coy, S., Rubin, L. L., and McMahon, A. P. (2012) Selective identification of hedgehog pathway antagonists by direct analysis of smoothened ciliary translocation. ACS Chem Biol 7, 1040–1048

