## Supplementary Figure and Table Legends for "Estrogen-related receptor beta activation and isoform shifting by cdc2-like kinase inhibition restricts migration and intracranial tumor growth in glioblastoma"

**Figures**

**Supplementary Figure 1**. **(A)** ESRRB mRNA expression in human brain as compared to estrogen receptor α (ERα, gene = *ESR1*) and OLIG1, a key regulator of neural progenitor cells. **(B)** ERRβ2 and ERRβsf protein are expressed in immortalized human MO3.13 oligodendrocyte cells. **(C)** Immunofluorescent staining of ERRβ2 and ERRβsf depicts nuclear localization of ERRβsf, and nuclear and cytoplasmic localization of ERRβ2, in MO3.13 oligodendrocyte cells.

**Supplementary Figure 2**. Knockdown of ERRβ2 significantly enhances T98G cell migration, as measured by transwell migration assay. Data are presented as the median (line) ± minimum/maximum for 3-5 fields of view per transwell filer, in each of at least two independent biological replicates. Data were analyzed by Mann-Whitney test at each time point, where **** denotes p<0.0001.

**Supplementary Figure 3**. **(A)** Cortactin mRNA expression (gene = *CTTN*) increases with increasing severity of glioma. RNAseq data from the from the Chinese Glioma Genome Atlas, obtained through the International Cancer Genome Consortium, were plotted in R. A = astrocytoma, AA = anaplastic astrocytoma, AO = anaplastic oligodendroglioma, AOA = anaplastic oligoastrocytoma, O = oligodendroglioma, OA = oligoastrocytoma, GBM = glioblastoma, r = recurrent, s = secondary. **(B)** Cortactin mRNA expression is significantly increased at the invasive front of GBM. RNAseq expression Z scores from micro dissected tumors obtained from the Ivy Glioblastoma Atlas Project through the Allen Brain Institute were analyzed by Mann-Whitney test comparing the cellular tumor to the leading edge. * denotes p<0.05.

**Supplementary Figure 4**. **(A, B)** Expression and phosphorylation of SR proteins in 42MGBA-TMZres (A) and T98G cells (B) treated with 50 μM TG-003, the combination of 50 μM TG-003 and 5 μM DY131, or DMSO for 24 hours.

**Supplementary Figure 5**. Full cell cycle profile of immortalized oligodendrocyte MO3.13 and GBM cell lines treated with 5 μM DY131, 50 μM TG-003, the combination of 5 μM DY131 + 50 μM TG-003 (TGDY), or DMSO control for 24 hours. For each panel, data are presented as the mean ± SD for 3 independent biological replicates. For T98G cells, data show an identical magnitude of G2/M arrest caused by the combination of 10 μM DY131 + silencing of ERRβsf (adapted from ref. 19) as that seen in response to the combination of 5 μM DY131 + 50 μM TG-003 (TG/DY).

**Tables**

**Supplementary Table 1**. Clinical, molecular, and pathological data for PDX models from the Mayo Clinic Brain Tumor Patient-Derived Xenograft National Resource. Tab 1 shows molecular and pathological data, and tab 2 shows clinical data. Full data for all PDX models in the resource are available at .

**Supplementary Table 2.** ERRβ splice variant F domain analysis. Sequence of the extended carboxyl terminal F domains of ERRβ2 and ERRβ-Δ10 were analyzed using ScanSite and medium stringency settings.
