## Supplementary figures and images for "Estrogen-related receptor beta activation and isoform shifting by cdc2-like kinase inhibition restricts migration and intracranial tumor growth in glioblastoma"

### Supplementary Figure 1

Supplementary Figure 1

A

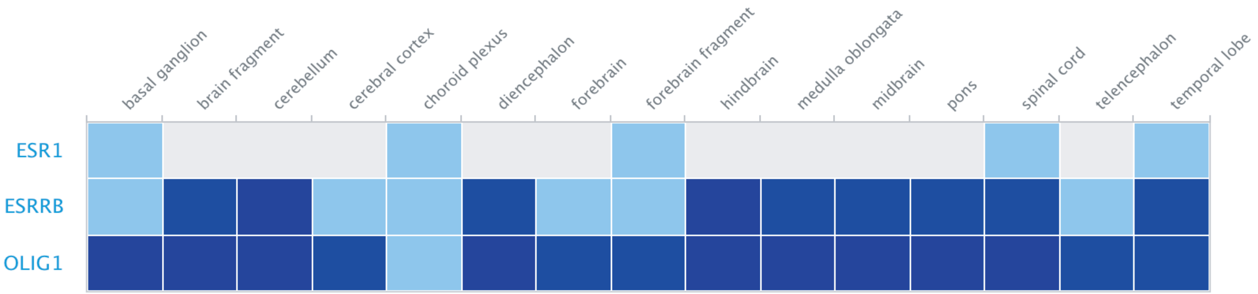


B

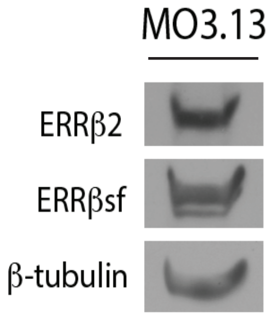


C

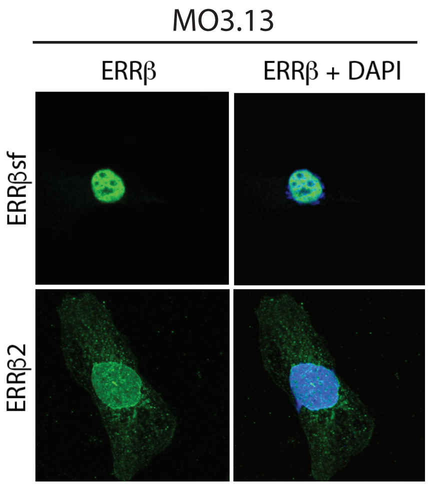

### Supplementary Figure 2

Supplementary Figure 2

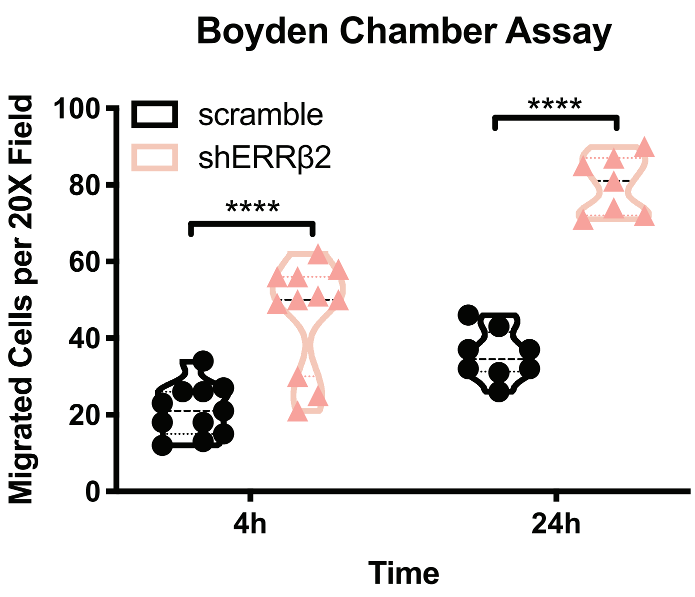

### Supplementary Figure 3

Supplementary Figure 3

A

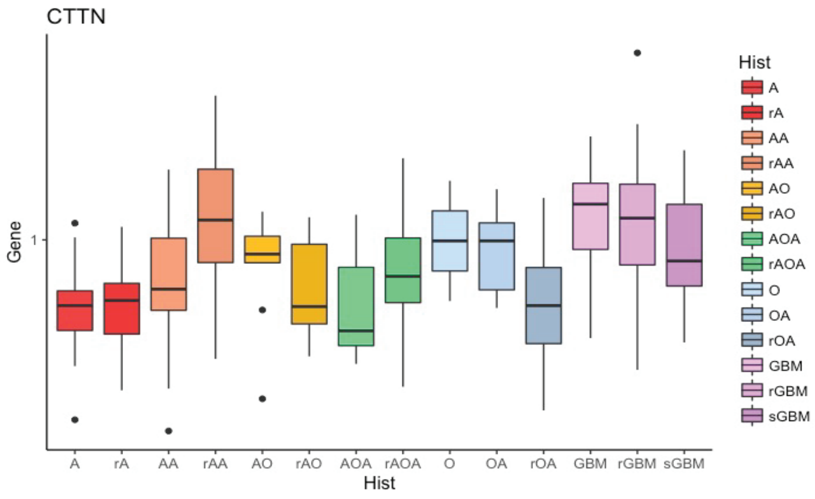


B

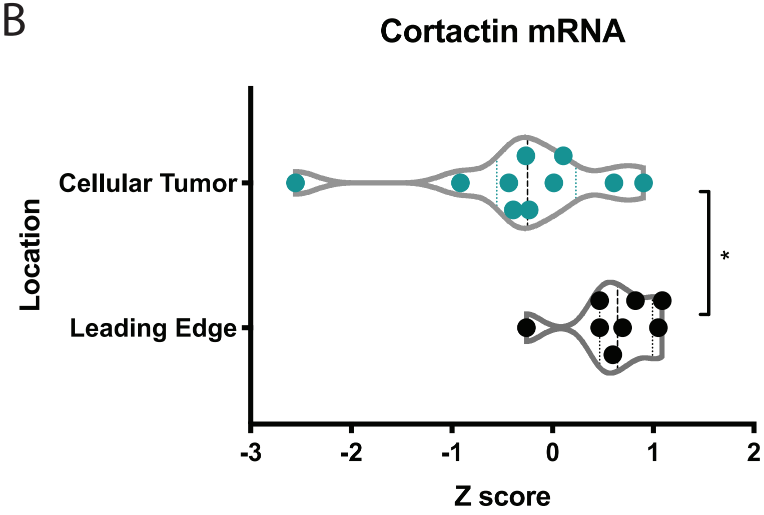

### Supplementary Figure 4

Supplementary Figure 4

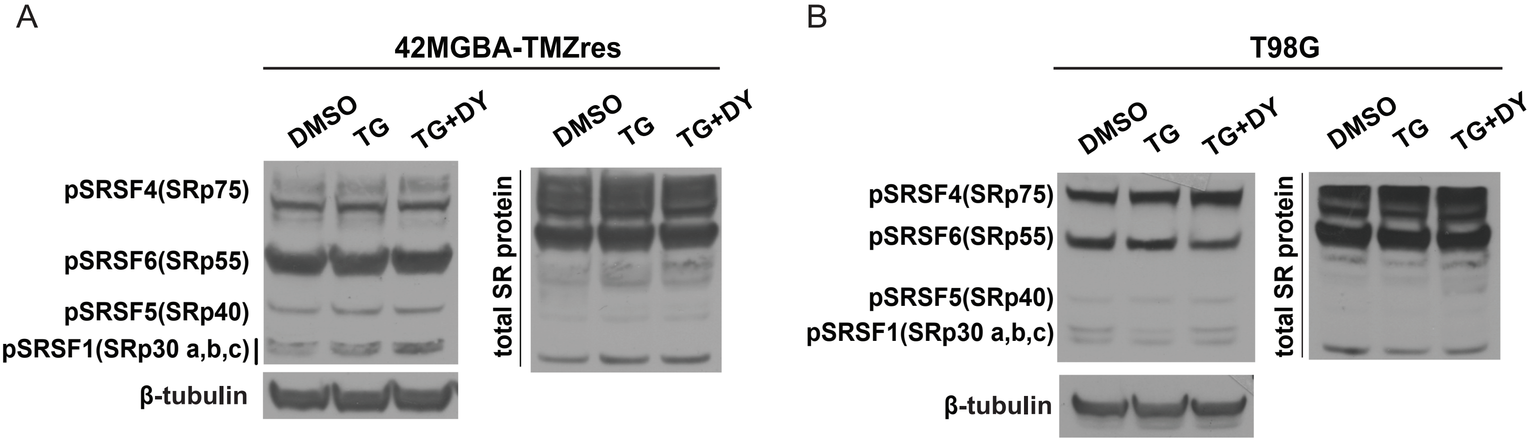

### Supplementary Figure 5

#### Supplementary Figure 5

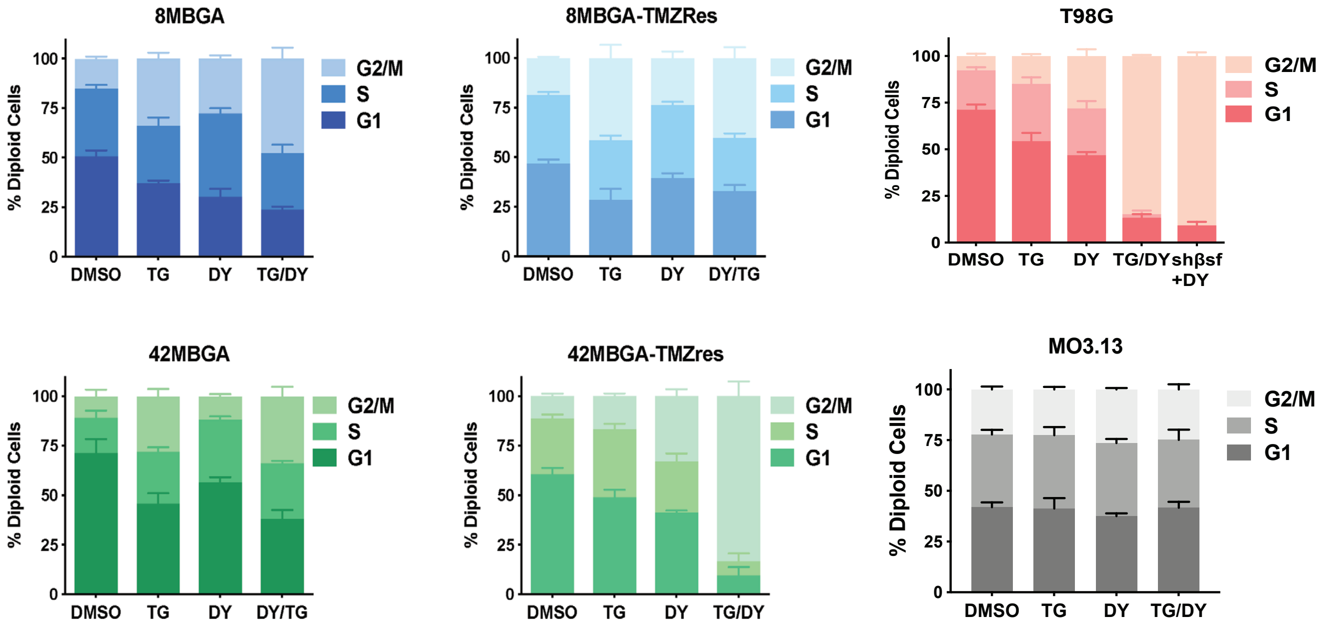
