## Supplementary Table 1 for "Estrogen-related receptor beta activation and isoform shifting by cdc2-like kinase inhibition restricts migration and intracranial tumor growth in glioblastoma"

### Supplementary Table 1 - Mayo PDX Characteristics

| GBM | Xenograft start | P, RC, or 2° XG IMPLANT | HUMAN SUBJECT DIAGNOSIS | SEX |
|---|---|---|---|---|
| 6 | 2000 | PRIMARY | GBM | M |
| 8 | 2000 | PRIMARY | GBM | F |
| 10 | 2001 | RECURRENT | OA | M |
| 12 | 2001 | PRIMARY | GBM | M |
| 14 | 2001 | RECURRENT | GBM | M |
| 22 | 2002 | PRIMARY | GS | M |
| 26 | 2002 | PRIMARY | GBM | M |
| 28 | 2002 | PRIMARY | GS | M |
| 38 | 2003 | PRIMARY | GBM | F |
| 39 | 2003 | PRIMARY | GBM | M |
| 43 | 2003 | PRIMARY | GBM | M |
| 44 | 2003 | PRIMARY | GBM | F |
| 46 | 2003 | RECURRENT | GBM | M |
| 59 | 2004 | PRIMARY | GBM | F |
| 64 | 2006 | RECURRENT | GBM | F |

| | |
|---|---|
| # Primary | 11 |
| # Secondary | 0 |
| # Recurrent | 4 |
| total | 15 |

| Defined from Methylomics | Defined by 250K Array CgH | | Defined by Sarkaria MS-PCR |
|---|---|---|---|
| **GBM MOLECULAR SUBTYPE** | **EGFR AMP.** | **EGFR MUT.** | **MGMT METHYLATION** |
| C | Y | VIII | U |
| C | Y | | M |
| NA | N | | U |
| C | Y | | M |
| P | N | | U |
| C | N | | M |
| NA | Y | | U |
| C | N | | U |
| P | Y | | U |
| C | Y | VIII | M |
| NA | N | | U |
| NA | N | | U |
| C | Y | VIII | M |
| C | Y | VIII | M |
| P | Y | | U |

| | |
|---|---|
| # methylated | 6 |
| # unmethylated | 9 |
| | 15 |

| Defined by MDAnderson CLIA MS-PCR | Defined by Sanger Sequencing | | |
|---|---|---|---|
| **MGMT METHYLATION2** | **TERT** | **IDH1** | **IDH2** |
| Indeterminate | C228T | wt | wt |
| M | C228T | wt | wt |
| U | C228T | wt | wt |
| M | C250T | wt | wt |
| U | C228T | wt | wt |
| M | C250T | wt | wt |
| U | C228T | wt | wt |
| U | C228T | wt | wt |
| U | C228T | wt | wt |
| M | C250T | wt | wt |
| M | C228T | wt | wt |
| U | C228T | wt | wt |
| M | C228T | wt | wt |
| M | C228T | wt | wt |
| M | C228T | wt | wt |

0
0

### Supplementary Table 1 - Mayo PDX Characteristics

| GBM | GBM. | SEX | age at dx | Yearn of DEATH or LAST F/U | STATUS 0=DEAD 1=ALIVE 2=LTF |
|---|---|---|---|---|---|
| 6 | 6 | M | 65 | 2001 | 0 |
| 8 | 8 | F | 75 | 2002 | 0 |
| 10 | 10 | M | 41 | 2001 | 0 |
| 12 | 12 | M | 69 | 2001 | 0 |
| 14 | 14 | M | 58 | 2002 | 0 |
| 22 | 22 | M | 80 | 2002 | 0 |
| 26 | 26 | M | 49 | 2003 | 0 |
| 28 | 28 | M | 68 | 2003 | 0 |
| 38 | 38 | F | 72 | 2005 | 0 |
| 39 | 39 | M | 51 | 2006 | 0 |
| 43 | 43 | M | 69 | 2004 | 0 |
| 44 | 44 | F | 80 | 2005 | 0 |
| 46 | 46 | M | 56 | 2004 | 0 |
| 59 | 59 | F | 83 | 2005 | 0 |
| 64 | 64 | F | 64 | 2006 | 0 |

| SURGERY/BIOPSY Year | GRADE (INITIAL) | HISTOLOGY | Year XG IMPLANTED |
|---|---|---|---|
| 2000 | 4 | Fibrillary Astrocytoma | 2000 |
| 2000 | 4 | Fibrillary Astrocytoma | 2000 |
| 1999 | 4 | Oligoastrocytoma | 2001 |
| 2001 | 4 | Glioblastoma Multiforme | 2001 |
| 2001 | 4 | Fibrillary Astrocytoma | 2001 |
| 2002 | 4 | Gliosarcoma | 2002 |
| 2002 | 4 | Fibrillary Astrocytoma | 2002 |
| 2002 | 4 | Gliosarcoma | 2002 |
| 2003 | 4 | Fibrillary Gemistocytic Astrocytoma | 2003 |
| 2003 | 4 | Small Cell Astrocytoma | 2003 |
| 2003 | 4 | Fibrillary Astrocytoma | 2003 |
| 2003 | 4 | Fibrillary Astrocytoma | 2003 |
| 2002 | 4 | Fibrillary Astrocytoma | 2003 |
| 2004 | 4 | Fibrillary & Gemistocytic Astrocytoma | 2004 |
| 2005 | 4 | Fibrillary Astrocytoma | 2006 |

| THERAPIES PRIOR TO XG *(NPT = no prior tx)* | OS from Dx | OS from xenograft (yrs) | TYPE OF XG IMPLANT 1=primary 2=recurrent 3=secondary |
|---|---|---|---|
| NPT | 1.05 | 1.05 | 1 |
| NPT | 1.34 | 1.34 | 1 |
| BCNU, Thalidomide, Carboplatin, TMZ | 2.50 | 0.62 | 2 |
| NPT | 0.24 | 0.24 | 1 |
| RT, Gefitinib | 0.90 | 0.38 | 2 |
| NPT | 0.21 | 0.21 | 1 |
| NPT | 0.73 | 0.73 | 1 |
| NPT | 0.67 | 0.50 | 1 |
| NPT | 1.38 | 1.38 | 1 |
| NPT | 2.41 | 2.41 | 1 |
| NPT | 0.26 | 0.26 | 1 |
| NPT | 1.40 | 1.36 | 1 |
| OSI-774, RT, BCNU, TMZ (x1 cycle) | 1.56 | 0.50 | 2 |
| NPT | 1.03 | 1.03 | 1 |
| OSI-774, RT, TMZ | 1.02 | 0.13 | 2 |

| GRADE (IMPLANT) | CLINICAL MGMT (≥ 2=METHYLATED) | 1ST TX Year | 1ST TX REGIMEN | PFS #1 (WEEKS) |
|---|---|---|---|---|
| 4 | - | 2000 | RT | 26.86 |
| 4 | - | 2001 | RT | 33.86 |
| 4 | - | 1999 | Observation | 19.14 |
| 4 | - | 2001 | RT | |
| 4 | - | 2001 | RT + Gefitinib | 23.29 |
| 4 | - | - | - | |
| 4 | - | 2002 | RT | 22.57 |
| 4 | - | 2002 | RT | |
| 4 | - | 2003 | RT | 13.43 |
| 4 | - | 2003 | RT | 25.71 |
| 4 | - | 2003 | RT | 4.57 |
| 4 | - | - | Palliative | |
| 4 | - | 2002 | OSI-774/RT | 25.14 |
| 4 | - | 2004 | RT | |
| 4 | - | 2005 | OSI-774/RT/TMZ + OSI-774/TMZ | 41.57 |

| CLINICAL TRIAL # | 1ST FAILURE Year | 1ST FAILURE REGIMEN | PFS #2 (WEEKS) | 2ND FAILURE Year |
|---|---|---|---|---|
| - | 2001 | BCNU | 17.29 | 2001 |
| - | 2001 | Gamma Knife | | - |
| - | 1999 | Craniotomy/BCNU Wafers + RT | 40.86 | 2000 |
| - | - | - | | - |
| N0074 | 2001 | Craniotomy + BCNU | 15.00 | 2002 |
| - | - | - | | - |
| - | 2003 | TMZ | | - |
| - | - | - | | - |
| - | 2004 | TMZ | 5.00 | 2004 |
| - | 2004 | TMZ | 38.14 | 2005 |
| - | 2004 | Palliative | | - |
| - | - | - | | - |
| N0177 (S2) | 2003 | BCNU | 20.29 | 2003 |
| - | - | - | | - |
| N0177 (S3) | 2006 | Craniotomy | | - |

| 2ND FAILURE REGIMEN | PFS #3 (WEEKS) | 3RD FAILURE Year | 3RD FAILURE REGIMEN |
|---|---|---|---|
| Continued BCNU | | - | - |
| - | | - | - |
| Thalidomide + 1D Carboplatin | 26.14 | 2000 | TMZ |
| - | | - | - |
| - | | - | - |
| - | | - | - |
| - | | - | - |
| - | | - | - |
| Temsirolimus | 28.57 | 2004 | BCNU |
| Craniotomy + BCNU | | - | - |
| - | | - | - |
| - | | - | - |
| TMZ | 5.57 | 2003 | Craniotomy + Irinotecan |
| - | | - | - |
| - | | - | - |

| PFS #4 (WEEKS) | 4TH FAILURE Year | 4TH FAILURE REGIMEN |
|---|---|---|
| | - | - |
| | - | - |
| 10.71 | 2001 | Craniotomy + Irinotecan |
| | - | - |
| | - | - |
| | - | - |
| | - | - |
| | - | - |
| | - | - |
| | - | - |
| | - | - |
| | - | - |
| 13.14 | 2004 | Etoposide |
| | - | - |
| | - | - |

| NOTES |
| --- |
| Pt. expired elsewhere |
| Pt's oncological tx is complicated by multiple infections |
| Prior to initiation of RT, MRI scans showed progression of disease & pt opted not to pursue therapy |
| Hx of plasmacytoma for which he underwent blood/marrow transplant 8/17/99 |
