## Supplementary Table 2 for "Estrogen-related receptor beta activation and isoform shifting by cdc2-like kinase inhibition restricts migration and intracranial tumor growth in glioblastoma"

**Supplementary Table 2 - ERRβ F Domain Motif Analysis**

**ERRβ2**

| MOTIF_NAME | MOTIF_GROUP | SCORE | PERCENTILE | PROTEIN | SITE | SITE_SEQUENCE | SURFACE_ACCESS_VALUE |
|---|---|---|---|---|---|---|---|
| Cdk5 Kinase | Proline-dependent serine/threonine kinase group | 0.4372 | 0.005277516 | beta2 AF2 | S19 | RPLEQVPsPLHRATK | 0.6115824 |
| Cdc2 Kinase | Proline-dependent serine/threonine kinase group | 0.5091 | 0.008282387 | beta2 AF2 | S19 | RPLEQVPsPLHRATK | 0.6115824 |
| Erk D-domain | Kinase binding site group | 0.5793 | 0.006573512 | beta2 AF2 | L33 | KRQHVHFITPLPPPP | 0.40981248 |
| Erk1 Kinase | Proline-dependent serine/threonine kinase group | 0.4059 | 0.001022549 | beta2 AF2 | T34 | RQHVHFLtPLPPPPS | 0.465696 |
| Cortactin SH3 | Src homology 3 group | 0.574 | 0.008602443 | beta2 AF2 | P37 | VHFLTPLpPPPSVAW | 1.670625 |
| Src SH3 | Src homology 3 group | 0.4637 | 0.006133115 | beta2 AF2 | P37 | VHFLTPLpPPPSVAW | 1.670625 |
| AMP_Kinase | Basophilic serine/threonine kinase | 0.5479 | 0.002540861 | beta2 AF2 | S41 | TPLPPPPsVAWVGTA | 0.57891834 |

**ERRβsf**

| MOTIF_NAME | MOTIF_GROUP | SCORE | PERCENTILE | PROTEIN | SITE | SITE_SEQUENCE | SURFACE_ACCESS_VALUE |
|---|---|---|---|---|---|---|---|
| Cdk5 Kinase | Proline-dependent serine/threonine kinase group | 0.4605 | 0.007251541 | delta10 AF2 | S8 | GQEQLRGsPKDERMS | 3.074035536 |
| Amphiphysin SH3 | Src homology 3 group | 0.5257 | 0.004216721 | delta10 AF2 | P37 | SRDQSNSpGIPNPRP | 0.85065552 |
| CDK1 motif 2 - [ST]PxxK | Proline-dependent serine/threonine kinase group | 0.4364 | 0.009011628 | delta10 AF2 | S46 | IPNPRPSsPTPLNER | 2.19594375 |
| Erk1 Kinase | Proline-dependent serine/threonine kinase group | 0.4776 | 0.004177797 | delta10 AF2 | S46 | IPNPRPSsPTPLNER | 2.19594375 |
| CDK1 motif 2 - [ST]PxxK | Proline-dependent serine/threonine kinase group | 0.39 | 0.004659988 | delta10 AF2 | S58 | NERGRQIsPSTRTPG | 1.11495384 |
| CDK1 motif 1 - [ST]Px[KR]x | Proline-dependent serine/threonine kinase group | 0.4227 | 0.007863007 | delta10 AF2 | S58 | NERGRQIsPSTRTPG | 1.11495384 |
| Calmodulin dependent Kinase 2 | Basophilic serine/threonine kinase | 0.4693 | 0.005202412 | delta10 AF2 | S58 | NERGRQIsPSTRTPG | 1.11495384 |
| Cdk5 Kinase | Proline-dependent serine/threonine kinase group | 0.4788 | 0.009183102 | delta10 AF2 | S58 | NERGRQIsPSTRTPG | 1.11495384 |
| Cdc2 Kinase | Proline-dependent serine/threonine kinase group | 0.4816 | 0.005482309 | delta10 AF2 | S58 | NERGRQIsPSTRTPG | 1.11495384 |
| Akt Kinase | Basophilic serine/threonine kinase | 0.5404 | 0.005897885 | delta10 AF2 | S58 | NERGRQIsPSTRTPG | 1.11495384 |
| CDK1 motif 2 - [ST]PxxK | Proline-dependent serine/threonine kinase group | 0.3887 | 0.004565373 | delta10 AF2 | T63 | QISPSTRtPGGQGKH | 1.41571584 |
| CDK1 motif 1 - [ST]Px[KR]x | Proline-dependent serine/threonine kinase group | 0.4216 | 0.007782861 | delta10 AF2 | T63 | QISPSTRtPGGQGKH | 1.41571584 |
